# Pathogenicity of autoreactive T_H_17 cells is modulated by the TRPM7 kinase

**DOI:** 10.64898/2026.10.27.762170

**Authors:** Katharina Jacob, Anna Sophie Thomann, Anna Madlmayr, Katharina Horner, Neysa Rodriguez Murillo, Dorothea Lewitz, Franz Bauernschmitt, Lisa Ann Gerdes, Christian Weber, Hendrik Schulze-Koops, Masayuki Matsushita, Thomas Gudermann, Birgit Karner-Hoeger, Anneli Peters, Susanna Zierler

**Affiliations:** Walther Straub Institute of Pharmacology and Toxicology, Ludwig-Maximilians Universität München, Munich, Germany; Institute for Cardiovascular Prevention (IPEK), Ludwig-Maximilians Universität München, Munich, Germany; Institute of Clinical Neuroimmunology, University Hospital and Biomedical Center, Ludwig-Maximilians Universität München, Munich, Germany; Institute of Pharmacology, Faculty of Medicine, Johannes Kepler University Linz, Linz, Austria; Munich Cluster of Systems Neurology (SyNergy), Munich, Germany; Division of Rheumatology and Clinical Immunology, Department of Medicine IV, LMU University Hospital, Ludwig-Maximilians Universität München, Munich, Germany; Department of Molecular and Cellular Physiology, Graduate School of Medicine, University of the Ryukyus, Okinawa, Japan; Clinical Research Institute for Inflammation Medicine, Johannes Kepler University Linz, Linz, Austria

## Abstract

The melastatin-like *transient receptor potential* channel, TRPM7, is crucial for signaling in leukocytes. It harbors a channel domain, permeable to divalent cations as well as a serine/threonine α-kinase, phosphorylating *inter alia* SMAD2. TRPM7 kinase activity is essential for the differentiation of non-pathogenic T helper 17 (T_H_17) cells as well as for T cell colonization and alloreactivity in the gut. As pathogenic T_H_17 (pT_H_17) cells drive autoimmune diseases, we studied TRPM7 kinase involvement in pT_H_17 polarization and function as well as in the development of multiple sclerosis (MS), using human T cells and established murine experimental autoimmune encephalomyelitis (EAE) models. Our results reveal an elevated TRPM7 activity in T effector cells derived from the cerebrospinal fluid (CSF) of MS patients, in line with higher functional TRPM7 expression on human T_H_17 cells. Utilizing a mouse line with disrupted kinase activity (*Trpm7^R/R^*), we uncovered both impaired differentiation of pathogenic T_H_17 cells as well as a significantly milder course of T_H_17-mediated transfer EAE in *Trpm7^R/R^* mice compared to *Trpm7^+/+^* controls. In the CNS of *Trpm7^R/R^* mice, the number of FOXP3^+^ T cells (T_regs_) was increased, while fewer T_H_17 cells infiltrated the CNS due to a migration defect of transferred pT_H_17 cells in the *Trpm7^R/R^* hosts and a failure to reactivate in the intestine, due to reduced MHCII signaling. Thus, inactivation of TRPM7 kinase may represent a promising therapeutic strategy targeting both differentiation and reactivation of pT_H_17 cells while preserving T_regs_ function and thereby dampening CNS inflammation in MS.

**Graphical Abstract:** 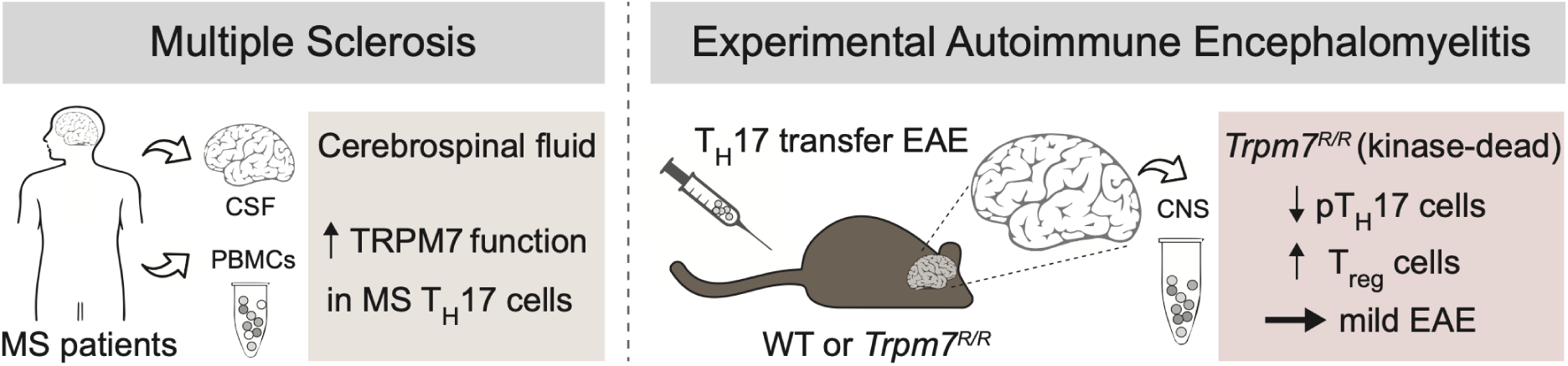

## Introduction

The *transient receptor potential* cation channel subfamily M, member 7 (TRPM7) is a cation channel with a C-terminally fused serine/threonine α-kinase (Cahalan, 2001; Nadler et al., 2001; Runnels et al., 2001; Ryazanova et al., 2004; Schmitz et al., 2003). It conducts divalent cations including calcium (Ca^2+^), magnesium (Mg^2+^) and zinc (Zn^2+^) and is an important component of cellular and systemic ion homeostasis (Li et al., 2007; Mittermeier et al., 2019; Monteilh-Zoller et al., 2003; Ryazanova et al., 2010; Schmitz et al., 2003). TRPM7 has been associated with numerous cellular functions including proliferation, differentiation, activation, and survival (McNeill et al., 2007; Romagnani et al., 2017; Sahni and Scharenberg, 2008; Wei et al., 2009; Wykes et al., 2007; Zierler et al., 2016; Zierler et al., 2011). In addition, it is strongly linked to immune cell development and homeostasis, and functions as an important signaling molecule particularly in lymphocytes (Krishnamoorthy et al., 2018; Madlmayr et al., 2026; Mellott et al., 2020; Mendu et al., 2020; Romagnani et al., 2017; Sahni and Scharenberg, 2008). Notably, inactivation of TRPM7 kinase activity does not affect channel function (Kaitsuka et al., 2014; Matsushita et al., 2005; Schmitz et al., 2003; Zierler et al., 2016), whereas the α-kinase requires the Mg^2+^ provided by the channel for its phosphotransferase activity (Matsushita et al., 2005; Runnels et al., 2001; Ryazanova et al., 2004).

In a previous study, we established a correlation between TRPM7 kinase activity and the development of graft *versus* host disease (GvHD) in mice (Romagnani et al., 2017). GvHD is a major complication after allogeneic blood stem cell transplantation(Akahoshi et al., 2024; Mehta and Koreth, 2025). Pro-inflammatory T helper 17 (T_H_17) cells differentiated from the transplanted donor cells turn against the host and cause serious and often lethal damage to organs such as the intestine (Harris and Karimi, 2023; Teshima et al., 2011). T_H_17 cells are also implicated in various autoimmune diseases, including psoriasis, rheumatoid arthritis, and multiple sclerosis (MS) (Goedken et al., 2025; Hedegaard et al., 2008; Hu et al., 2025; Kamada et al., 2013; Sarkar et al., 2025).

MS is the most common demyelinating disease in young adults with a range of disease courses, clinical symptoms and severities. Autoimmune inflammation manifests in the central nervous system (CNS), where the myelin sheath is targeted, leading to damage of the underlying axons and subsequent sensory impairments and paralysis (Jakimovski et al., 2024; Thompson et al., 2018). The exact etiology of MS is unclear but both genetic susceptibilities, as well as environmental factors including Vitamin D deficiency, smoking, composition of the intestinal microbiota, and viral infections with Epstein-Barr virus have been associated with the risk for disease development (Alfredsson and Olsson, 2019; Bjornevik et al., 2022; Correale et al., 2022; International Multiple Sclerosis Genetics, 2019). While the nature of the autoantigen is not clearly defined, autoreactive T_H_17 and T_H_1 T cell subtypes have been suggested to be main drivers of the inflammatory cascade (Comi et al., 2021; Hohlfeld et al., 2016). Recently, it has also been demonstrated that B cells are involved in the pathogenesis of MS (Comi et al., 2021; Hohlfeld et al., 2016).

In this study we investigated whether TRPM7 kinase plays a role in function and pathogenicity of autoreactive T_H_17 cells in CNS autoimmunity. We assessed functional TRPM7 expression in CSF cells of MS patients, T_H_17 and regulatory T cell (T_reg_) differentiation and function *ex vivo,* as well as *in vivo* in the kinase-deficient mouse model, *Trpm7^R/R^*, in two distinct animal models of MS: in myelin oligodendrocyte glycoprotein (MOG)-induced experimental autoimmune encephalomyelitis (EAE), and in T_H_17 adoptive transfer EAE. The results of the present study highlight the significance of the TRPM7 kinase as an important driver of CNS autoimmunity and as a promising pharmacologic target.

## Results

### Functional TRPM7 surface expression varies in distinct human T cell subsets

The channel-kinase TRPM7 has been shown to be particularly important for lymphocyte function, with channel conductance being essential for kinase activity. Recently, we demonstrated distinct effects of TRPM7 activity on human effector T cell polarization (Madlmayr et al., 2026). As the kinase moiety of TRPM7 has been implicated in driving inflammatory conditions and T_H_17 polarization in mice, we were wondering whether it was also instrumental for T_H_17-driven autoimmune diseases, such as MS. To understand whether changes in functional expression of TRPM7 may be associated with disease, we tested TRPM7 channel activity in cells isolated from untreated, newly diagnosed MS patients. To determine TRPM7 activity on T effector cells (T_eff_) *ex vivo*, we isolated cells directly from the cerebrospinal fluid (CSF) of MS patients (n=3), which is enriched for effector CD4^+^ T cells (Beltran et al., 2019; Eschborn et al., 2021; Gross et al., 2021; Jacobs et al., 2025). We compared TRPM7 currents of these CSF-derived CD4^+^ T cells to CD4^+^ T cells obtained from peripheral blood mononuclear cells (PBMC) of the same patients (Fig. 1A-D). First, we confirmed characteristic TRPM7 current development and current-voltage (I/V) relationships after 300 s in Mg^2+^-free solutions, maximizing TRPM7 channel activity (Fig. 1B, C) using whole-cell patch-clamp. Utilizing these standard conditions, we identified that CD4^+^ T cells extracted from the CSF cells of MS patients exhibited larger TRPM7 currents compared to CD4^+^ T cells extracted from PBMCs of the same patients (Fig. 1 b-d). Thus, we hypothesized that the functional surface expression levels of TRPM7 at the plasma membrane (PM) of distinct effector CD4^+^ T cell subsets might vary. To address this, we isolated primary human naïve T cells, natural T helper 17 (nT_H_17) as well as natural regulatory T (nT_reg_) cells from healthy donors and measured TRPM7 channel activity utilizing established protocols (Ryazanova et al., 2010; Zierler et al., 2016) (Fig. 1E-L S6A, S7A). Again, we established characteristic TRPM7 current development and I/V relationships in these cells, after 300 s in Mg^2+^-free solutions (Fig. 1E). To estimate differences in functional TRPM7 channel expression at the cell surface, we applied a divalent-free solution (Fig. 1F; see methods), removing all inhibitory divalent cations of open TRPM7 channels and thus linearizing I/V-relationships (Fig. 1G, H). Utilizing this method, we determined the number of open channels per cell with the described single channel conductance (39 pS at -80mV) under these conditions (Kerschbaum and Cahalan, 1999; Kozak et al., 2002) (Fig. 1H, I). We calculated that nT_H_17 cells expressed significantly more functional TRPM7 channels at the PM (∼60 channels per cell) compared to nT_reg_ cells (∼30 channels per cell) or naïve T cells (∼40 channels per cell) (Fig. 1I).

**Figure 1.**
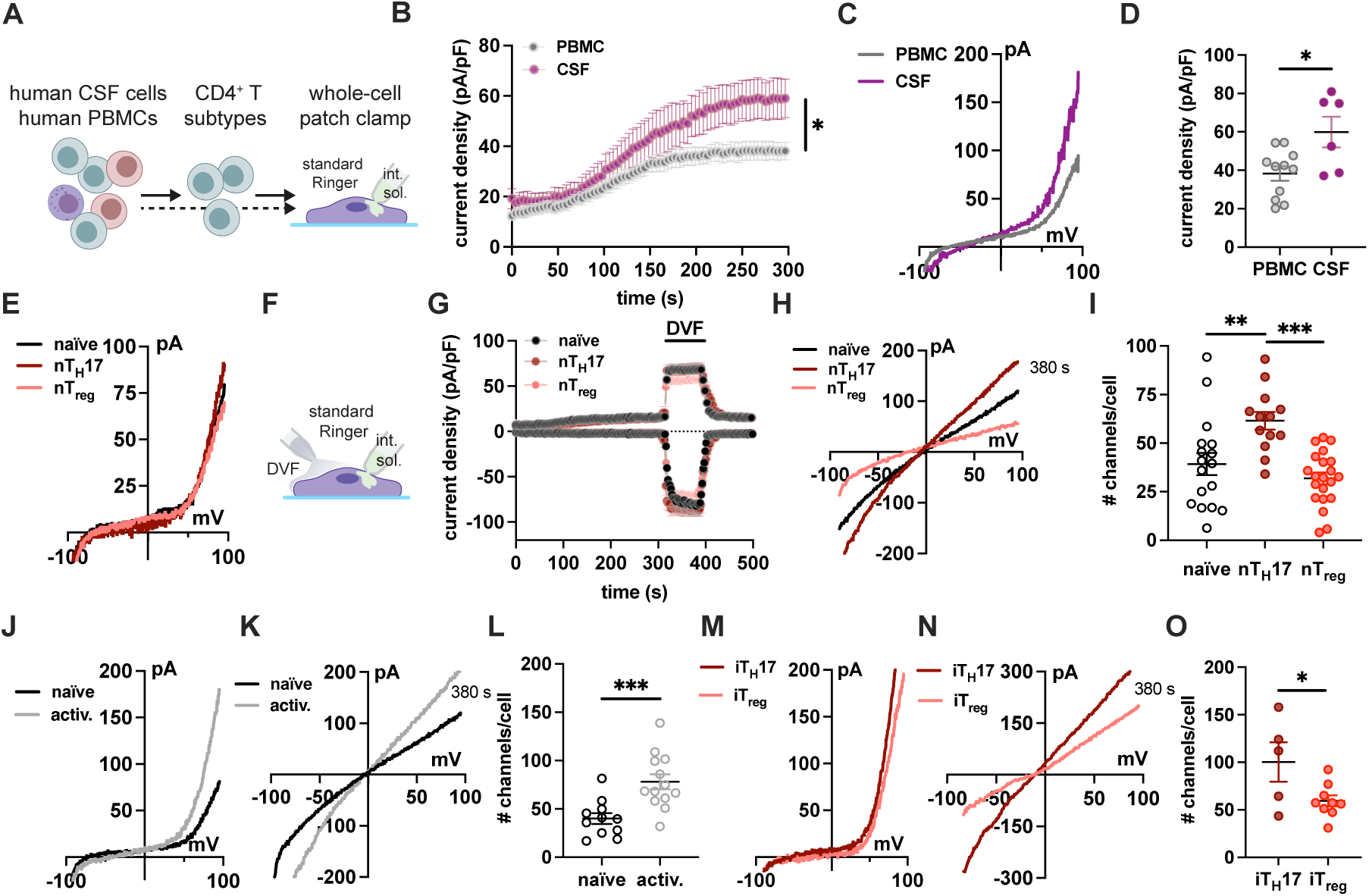
TRPM7 surface expression is upregulated in CD4^+^ T cells extracted from the CSF of MS patients as well as in human T_H_17 compared to T_reg_ cells. **(A)** Schematic drawing of the *ex vivo* immune cell preparation subjected to whole-cell patch-clamp technique. (B-D) Electrophysiological measurements of effector T cell populations isolated from peripheral blood (PBMC, gray) or CSF (purple) of MS patients. **(B)** Average TRPM7 current density, extracted at +80 mV was blotted *versus* time of the experiment in seconds (s). (**C** and **D**) Average current/voltage relationships (I/V) (C) and current density quantification of PBMC (purple, n=12 cells) and CSF cells (gray, n=6 cells) extracted at 300 s (D) from MS patients. (**E**) I/V-relationships of different T cell subpopulations isolated from PBMC of healthy donors; naïve T cells (black, n=11 cells), nT_H_17 (dark red, n=13) and nT_reg_ (light red, n=13 cells) extracted at 300 s. (**F**) Schematic illustration of the whole-cell patch clamp method with application of divalent-free (DVF) solution. (**G**) Average TRPM7 current densities, extracted at +80 mV, were blotted *versus* time of the experiment in seconds (s). (**H**) Average I/V relationships extracted at 380 s. (**I**) Calculated number of active TRPM7 channels/cell. (**J**-**L**) Average I/V relationships extracted at 300 s (J) and 380 s (K), as wells as the quantification of open TRPM7 channels/cell (L) for naïve T cells (black, n=11 cells) and PMA/ionomycin-activated T cells (gray, n=13 cells). (**M**-**O**) Representative I/V relationships extracted at 300 s (M) and 380 s (N), as well as the quantification of open TRPM7 channels/cell (O) for iT_H_17 (dark red, n=5 cells) and iT_reg_ (light red, n=9 cells) cells. Data are presented as mean ± SEM; *p < 0.05, **p < 0.01, ***p < 0.001, unpaired students t-test, except in L) one-way ANOVA was applied.

To verify whether TRPM7 channels are upregulated at the cell surface upon T cell activation, we stimulated human naïve T cells under supra-physiological conditions via PMA and ionomycin for 48h and measured characteristic TRPM7 I/V relationships, using standard conditions (Ryazanova et al., 2010; Zierler et al., 2016) (Fig. 1J-L). Already under standard patch-clamp conditions, we observed higher average current activities in activated T cells, compared to resting naïve T cells (Fig. 1J). When applying DVF solutions, these differences became more pronounced and the calculation of active channels per cell revealed a significant upregulation of TRPM7 at the PM upon T cell stimulation (∼80 *versus* ∼40 channels per cell, comparing activated and naïve cells, respectively) (Fig. 1K, L). Finally, we activated and polarized naïve human T cells into both T_H_17 cells and T_regs_ *in vitro*, using specific polarizing milieus and magnetic cell sorting (iT_reg_ cells) or a cytokine capture assay (Leipe et al., 2020a; Leipe et al., 2020b) (iT_H_17 cells), respectively. In line with what we observed in the primary nT_H_17 and nT_reg_ cells, we detected significantly higher numbers of active TRPM7 channels in human iT_H_17 (∼100 channels per cell) compared to iT_reg_ cells (∼55 channels per cell) (Fig. 1M-O). Thus, TRPM7 is highly expressed in proinflammatory effector T cell subsets but only to a much lower degree in non-activated and regulatory T cells.

### Molecular signature and transcriptome differences between TRPM7 kinase deficient and wild-type pathogenic T_H_17 and T_reg_ cells

The channel-kinase TRPM7 is important for lymphocyte function, with a functional channel moiety being indispensable for kinase activity (Madlmayr et al., 2026). As our results demonstrate an enhanced TRPM7 channel expression on human T_H_17 cells, we turned to our established mouse model, harboring a single point mutation at the active site of TRPM7 kinase (*Trpm7^R/R^*, p.1646R) to better characterize the effect of the kinase moiety (Romagnani et al., 2017). Previously, we demonstrated an impaired differentiation of T_H_17 cells upon interleukin (IL)-6 and transforming growth factor-β1 (TGF-β1) stimulation in TRPM7 kinase-deficient, *Trpm7^R/R^*, murine lymphocytes (Romagnani et al., 2017). Here, we asked whether TRPM7 kinase activity could be a major driver of the pathogenicity of T_H_17 cells and consequently, T_H_17-associated autoimmunity. Importantly, terminal differentiation and pathogenic effector functions of T_H_17 cells critically depend on the cytokine IL-23 and are further enhanced in an IL-1β-rich environment (Park and Ciofani, 2025; Peters et al., 2011).

Thus, we first differentiated naïve WT (*Trpm7^+/+^*) and *Trpm7^R/R^* CD4^+^ T cells using antibody-based T cell receptor (TCR) stimulation in a polarizing milieu containing the cytokines TGF-β1 and IL-6, as well as IL-23. To increase the pathogenicity of the T_H_17 cells even further, we added the inflammatory cytokine IL-1β (Fig. 2A) (Lee et al., 2012; Revu et al., 2018; Sutton et al., 2006). Compared to Trpm7^+/+^ T cells, significantly fewer *Trpm7^R/R^* T cells differentiated into pathogenic T_H_17 (pT_H_17) cells, in both the absence and presence of IL-1β (Fig. 2B).

**Figure 2.**
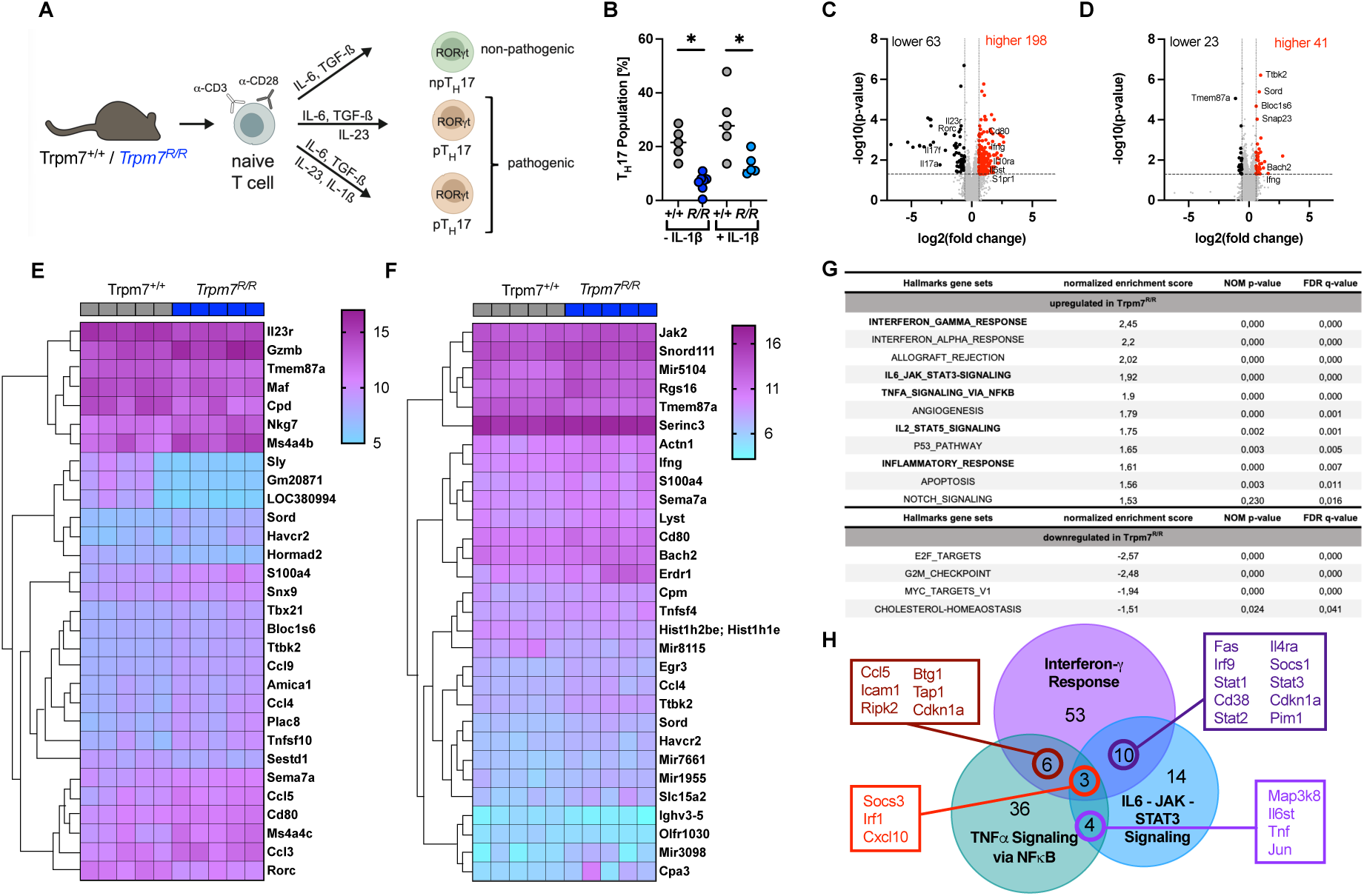
Genetic TRPM7 kinase inactivation inhibits murine pT_H_17 polarization and increases non-pathogenic but also inflammatory signatures. **(A)** T_H_17 cell polarization of naïve CD4^+^ T cells in distinct cytokine milieus leads to the differentiation of either non-pathogenic, or pathogenic T_H_17 (p T_H_17) cells. **(B)** Frequencies of differentiated T_H_17 cells after 4 days of culture quantified by flow cytometry. Data are shown as mean ± SEM (n = 5). Transcriptome analysis of pT_H_17 cells differentiated *in vitro* from naïve Trpm7^+/+^ (+/+) or *Trpm7^R/R^* (*R/R*) CD4^+^ T cells in the presence of IL-6, TGF-β1 and IL-23 with or without addition of IL-1β. **(C-D)** Volcano plot of differentially expressed genes (DEG) comparing Trpm7^+/+^ with *Trpm7^R/R^* in pT_H_17 cells differentiated without (C) and with (D) IL-1β. Black indicates down- and red up-regulated genes in *Trpm7^R/R^* compared to Trpm7^+/+^. **(E, F)** Heatmap illustrating the normalized gene expression levels of the top 30 altered genes between +/+ and *R/R* cells differentiated with indicated cytokines, (E) without IL-1β and (F) with IL-1β. **(G)** Hallmark gene sets, analyzed with GSEA. Gene sets related to inflammatory responses are shown in bold. **(H)** Venn diagram showing the overlapping genes between the Hallmark gene sets INTERFERON_GAMMA_RESPONSE, IL6&_JAK_STAT3_SIGNALING and TNFA_SIGNALING_VIA_NFKB \**p* < 0.05, \*\**p* < 0.01, \*\*\**p*< 0.001, ns, not significant, using multiple t-test, volcano plots and heatmaps were analyzed by the TAC software.

To gain insights into the molecular difference underlying TRPM7 kinase inactivation in pT_H_17 cell subsets, we analyzed gene expression patterns between differentially polarized Trpm7^+/+^ and *Trpm7^R/R^* T cells based on gene expression profiling microarray analysis (Fig. 2C-H). This transcriptome analysis revealed that, without IL-1β in the polarizing milieu, differences between *Trpm7^R/R^* and Trpm7^+/+^ pT_H_17 cells were more prominent (compare Fig. 2C and 2D) resulting in a total of 198 significantly up-regulated genes upon TRPM7 kinase inactivation (Fig. 2C). As illustrated in the volcano plot (Fig. 2C) as well as in the heatmap, the 30 most differentially expressed genes in *Trpm7^R/R^* compared to Trpm7^+/+^ pT_H_17 cells (Fig. 2E) include non-pathogenic T_H_17-signature genes such as *Il6st* (GP130), but also many immune-signaling (cytokine) genes (including *Ccl3* (MIP1-α), *Ccl4* (MIP1-β), *Ccl5* (RANTES) and *Ccl9* (MIP1-γ)), indicating that multiple genes involved in immunological pathways may be activated concordantly, possibly as a compensatory measure (Fig. 2C, E S1N, U,V). Thus, we performed gene-set enrichment analysis (GSEA) by using the MSigDB hallmark gene sets to determine whether specific gene sets were correlated with the inactivation of TRPM7 kinase. The GSEA enrichment analysis revealed that many immune-related gene sets were positively enriched in *Trpm7^R/R^* compared to Trpm7^+/+^ pT_H_17 cells, such as the gene set “HALLMARK_INTERFERON_GAMMA_RESPONSE”, whose expression is connected to genes up-regulated in response to interferon-γ. Notably, genes involved in the “HALLMARK_IL6_JAK_STAT3_SIGNALING” (genes up-regulated in response to the T_H_17 cell polarizing cytokine IL-6), the “HALLMARK_TNFA_SIGNALING_VIA_NFkB” (genes up-regulated in response TNFα and NFκB) and more general the “HALLMARK_INFLAMMATORY_RESPONSE” were also highly positively enriched in *Trpm7^R/R^* pT_H_17 cells (Fig. 2g-h, S1a-e). Visualizing overlapping genes for the top 3 upregulated Hallmark gene sets of interest with a Venn diagram, we identified *Socs3*, *Irf1* and *Cxcl10* as common significantly enriched genes (Fig. 2H; S1X, Y). *Socs3* encodes for the suppressor of cytokine signaling 3 and binds to active Janus kinases (e.g. JAK2) and specific cytokine receptors (e.g. gp130 and IL-6 receptor), thereby negatively regulating the JAK/STAT signaling pathway, controlling inflammation, immune cell differentiation, and metabolic processes (Carow and Rottenberg, 2014) and counteracting T_H_17 cell differentiation.(Qin et al., 2009) *Irf1* encodes for the transcription factor Interferon regulatory factor 1, which is activated by interferons (Wen et al., 2025) and viral infection, regulating innate and adaptive immunity as well as tumor suppression. (Zoler et al., 2026) *Cxcl10* encodes for C-X-C motif chemokine ligand 10 (CXCL10), also known as Interferon gamma-induced protein 10 (IP-10). Alterations in CXCL10 expression levels have been associated with inflammatory diseases including infectious diseases, immune dysfunction and tumor development. (Liu et al., 2011) In total, 59 genes were significantly downregulated upon TRPM7 kinase inactivation, including E2F target genes which are regulated by the E2F transcription factor family, primarily governing the G_1_-to-S phase cell cycle transition, DNA replication, and repair (Fig. 2G).

Analyzing specifically T_H_17-signature transcripts, a significant reduction of the average mRNA expression levels of the T_H_17-specific orphan nuclear receptor, *Rorc*, encoding for RORγt (Fig. 2C, E, S1F), as well as of the RORγt-dependent signature cytokines, *Il17a* and *Il17f* (Fig. 2C, E, S1G, P) was observed. Notably, upregulation of *Il23r,* which encodes for the IL-23 receptor (Fig. 2C, E, S1H) and has been directly associated with high pathogenicity (Miossec et al., 2009; Ramesh et al., 2014; Sallusto et al., 2012), was less severe upon TRPM7 kinase inhibition. Similarly, the average expression of the inflammatory chemokine receptor, *Ccr6*, which is induced during T_H_17 cell development by the synergistic actions of STAT3 and RORγt and also associated with autoimmune disease development (Miossec et al., 2009; Sallusto et al., 2012; Wang et al., 2009), was significantly reduced in *Trpm7^R/R^* compared to Trpm7^+/+^ pT_H_17 cells (Fig. S1J).

In the presence of IL-1β, only 40 genes were identified as significantly up-regulated (Fig. 2D), including the regulatory transcription factor BACH2, and only 23 genes as significantly down-regulated. Thus, most of the effects of genetic TRPM7 kinase inactivation on the expression of T_H_17-associated genes were abrogated by addition of IL-1β (Fig. S1 F-Y). Taken together, our results indicate that genetic inactivation of TRPM7 kinase promotes skewing towards a more anti-inflammatory non-pathogenic T_H_17 phenotype (Fig. S1).

As we observed increased expression of anti-inflammatory genes in *Trpm7^R/R^* compared to *Trpm7^+/+^* pT_H_17 cells, we also analyzed potential differences in regulatory T cell (T_reg_) subsets. Here, we first compared the relative number of natural regulatory T cells (nT_reg_), isolated from Trpm7^+/+^ and *Trpm7^R/R^* mice, and found that it was not affected by the loss of TRPM7 kinase activity (Fig. S2). Next, we differentiated naïve Trpm7^+/+^ and *Trpm7^R/R^* T cells into induced regulatory T cells (iT_reg_) *in vitro* using antibody-based TCR stimulation and a polarizing milieu containing TGF-β1 and IL-2. We found the frequency of *Foxp3*^+^ *Trpm7^R/R^* iT_reg_ to be slightly increased compared to *Trpm7^+/+^* controls (Fig. S2A), albeit not significantly. Analyzing average expression levels of anti-inflammatory mediators (Roychoudhuri et al., 2013; Sumida et al., 2024; Tang and Bluestone, 2008), no significant differences in mRNA expression of pro- or anti-inflammatory markers (Fig. S2 B-I) were detectable. Only the average *Il2* expression level was significantly higher in *Trpm7^R/R^* naïve T cells, compared to controls, but this effect was lost in the nT_reg_ population or upon differentiation into iT_reg_ (Fig. S2E), indicating that the regulatory T cell linage was only mildly affected by the loss of TRPM7 kinase activity.

### Loss of TRPM7 kinase activity is compensated for in EAE induced by immunization and adjuvant

Since we found modulation of gene expression in pT_H_17 cells by TRPM7 kinase as well as increased expression of TRPM7 in human T_H_17 cells *in vitro* and in CD4^+^ T_eff_ cells from MS patients *ex vivo*, we next investigated the role of TRPM7 kinase in T cell-driven autoimmunity in murine models of MS, termed experimental autoimmune encephalomyelitis (EAE). We first employed the conventional EAE model, in which disease is induced via immunization with myelin oligodendrocyte glycoprotein (MOG). To exclude differences between *Trpm7^+/+^* and *Trpm7^R/R^* mice under homeostatic conditions, we analyzed the distribution of different immune cell types in the spleen (SP) and central nervous system (CNS) and detected no significant alterations (Fig. S3 A-F). Next, we induced EAE via immunization with MOG_35-55_ emulsified in Complete Freund’s Adjuvant (CFA) plus two injections of pertussis toxin (PTX) (Fig. 3A) and compared disease development in *Trpm7^+/+^* and *Trpm7^R/R^* mice. Surprisingly, we did not observe any differences in disease incidence (Fig. 3B), weight loss (Fig. 3C, D) or disease score (Fig. E, F) between *Trpm7^+/+^* and *Trpm7^R/R^* mice. In line with this, the frequency of CNS-infiltrating CD4^+^ T cells was comparable between *Trpm7^+/+^* and *Trpm7^R/R^* mice (Fig. 3G) and both the frequency of FOXP3^+^ T_regs_ (Fig. 3H), as well as the frequency of IL-17^+^, IFNγ^+^ and IL-17^+^ IFNγ^+^ effector T cells (Fig. 3I) did not differ between the two genotypes. Similarly, there were no differences in overall CD4^+^ T - cell numbers, T_regs_ or cytokine-producing effector T cells in the spleens of these mice (Fig. S4A), indicating that neither T cell activation and differentiation nor CNS infiltration depends on TRPM7 kinase activity during this actively induced EAE model. Since PTX has been shown to affect Ca^2+^ influx in different settings (Tang et al., 2015) and is a known blocker of G-Protein-coupled receptors (GPCRs), which were shown to be capable of modulating TRPM7 channel activity (Takezawa et al., 2004), we excluded an impact of the administered PTX on TRPM7 channel function (Fig. S4 B, C). Taken together, we found that TRPM7 kinase is not required for EAE development after immunization. This may be explained by our findings that even though less TRPM7 kinase-deficient cells differentiate into T_H_17 cells, their inflammatory signature is largely normal in the presence of IL-1β (Fig. S1F-Y). Importantly, immunization with CFA + PTX induces a strong innate immune response including high levels of IL-1β (Benahmed et al., 2014; Shenderov et al., 2013) and might therefore override the defect of *Trpm7^R/R^* cells in pT_H_17 differentiation.

**Figure 3.**
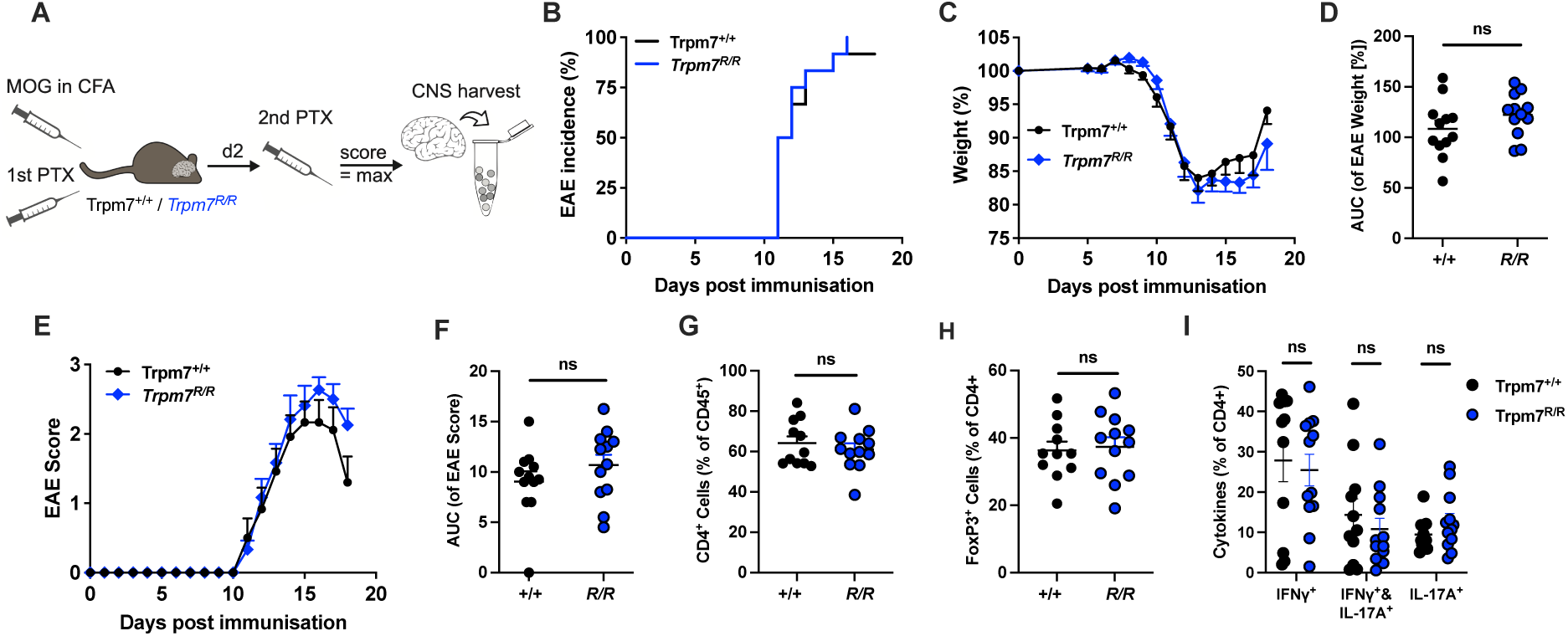
TRPM7 kinase activity is not required for EAE development after immunization with MOG/CFA and PTX. (**A**) Schematic experimental outline. Trpm7^+/+^ and *Trpm7^R/R^* mice were immunized by subcutaneous injection of MOG_35-55_ peptide emulsified in complete Freund’s adjuvant (CFA) containing *M. tuberculosis* and received two intraperitoneal injections of pertussis toxin (PTX) on days 0 and 2 after immunization. Mice were monitored for clinical signs of disease, and the CNS was harvested at the peak of the disease. (**B**-**D**) EAE incidence (B), body weight change over time (C) and body weight change quantified as area under the curve (AUC) (D). (**E**-**F**) EAE scores over time (E) and quantified as area under the curve (AUC) (F). Clinical data in B-F is shown as mean + SEM for 12 mice per group pooled from two independent experiments. (**G**) Frequency of CD4^+^ cells within the CD45^+^ population in the CNS of sick mice. (**H**-**I**) Frequency of FOXP3^+^ cells (H) and frequencies of different cytokine-producing cells (I) within the CD4^+^ population in the CNS of sick mice. Flow cytometry data in G-I is shown as mean + SEM for 11 (Trpm7^+/+^) or 12 sick mice (*Trpm7^R/R^*) per group. ns, not significant.

### TRPM7 kinase activity is crucial for disease development in T_H_17-mediated transfer EAE

It was shown that high levels of IL-1β are triggered by the adjuvant (CFA and PTX) used for immunization in the actively induced EAE model (Ronchi et al., 2016) and this may override the requirement for TRPM7 kinase in the function of pT_H_17 cells. However, such a strong stimulation of the innate immune response as induced in this setting, is unlikely to occur in human MS during priming of the autoreactive effector T cells. In addition, when patients are first diagnosed, autoreactive effector T cells are already present and may undergo reactivation events during clinical relapses (Afzali and Korn, 2025; Bielekova et al., 2000). Thus, we aimed to investigate whether already existing pT_H_17 cells could be modulated by a TRPM7 kinase-deficient environment and performed a T_H_17 adoptive transfer EAE, in which *in vitro* differentiated MOG-specific T_H_17 cells induce disease in an adjuvant-free setting (Jager et al., 2009) (Fig. 4A). Interestingly, we observed reduced EAE incidence (Fig. 4B) and severity (Fig. 4C-F) in TRPM7 kinase-deficient T_H_17 recipients as compared to *Trpm7^+/+^* control animals. This was associated with significantly reduced numbers of infiltrating CD4^+^ T cells in the CNS of sick *Trpm7^R/R^* mice at the peak of disease (Fig. 4G). Especially the frequency of transferred T_H_17 cells, identified by their transgenic T cell receptor chain Vα3.2, was strongly reduced in *Trpm7^R/R^* recipients compared to *Trpm7^+/+^* controls (Fig. 4H), whereas the frequency of FOXP3^+^ T_regs_ was increased in the CNS of *Trpm7^R/R^* mice (Fig.4I). In line with this, we detected an increased frequency of IFNγ^+^ CD4^+^ T cells in the CNS whereas the frequency of IL-17A^+^ CD4^+^ T cells was reduced in TRPM7 kinase-deficient recipients in the CNS and in the spleen (Fig. 4J, N). A similar reduction of Vα3.2^+^-transferred T_H_17 cells but no increase in T_regs_ was observed in the spleens of *Trpm7^R/R^* mice (Fig. 4K-M), indicative of impaired survival or reactivation of encephalitogenic T cells in the periphery of TRPM7 kinase-deficient hosts even prior to EAE onset.

**Figure 4.**
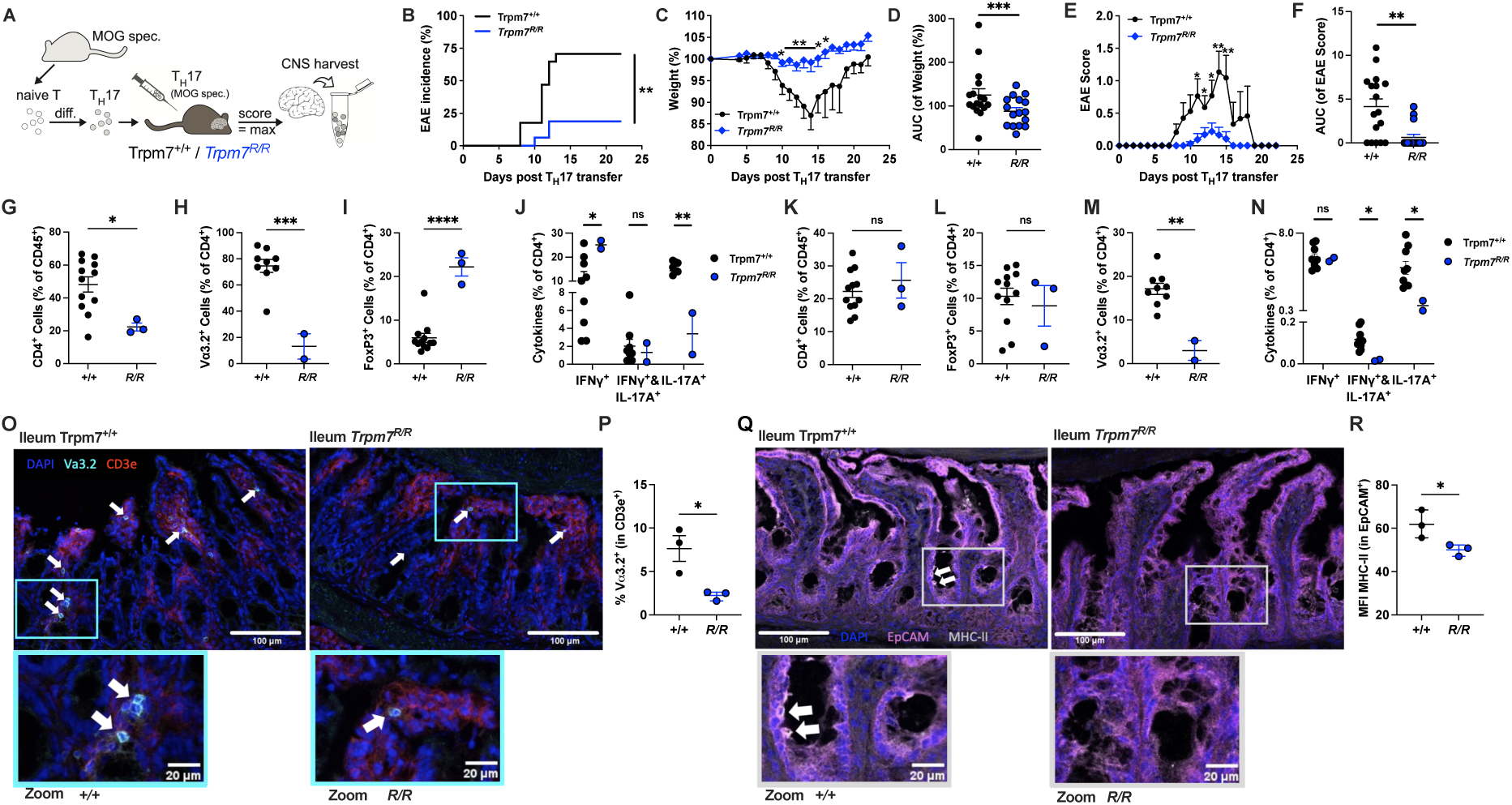
TRPM7 kinase activity is crucial for disease development in T_H_17-mediated transfer EAE. (**A**) Experimental outline. MOG-specific naïve CD4^+^ T cells isolated from transgenic 2D2 mice were differentiated into T_H_17 cells *in vitro* and injected intraperitoneally into Trpm7^+/+^ or *Trpm7^R/R^* recipients. Mice were monitored for clinical signs of disease; the CNS was harvested at the peak of disease. (**B**) EAE incidence, *\*\*p <* 0.01 Mantel-Cox Log-rank test. (**C**-**D**) Weight change over time (C) and quantified as area under the curve (AUC) (D). (**E**-**F**) EAE score over time (E) and quantified as area under the curve (AUC) (F). Clinical data in B-F is shown as mean ± SEM of 17 mice ( Trpm7^+/+^) or 16 mice *(Trpm7^R/R^)* per group pooled from 3 independent experiments. *p *<* 0.05, **p < 0.01, ***p < 0.001 unpaired t-test. (**G**) Frequency of CD4^+^ T cells within the CD45^+^ population in the CNS of sick mice. (**H**) Frequency of Vα3.2^+^ transferred T cells within the CD4^+^ population. (**I**-**J**) Frequency of FOXP3^+^ (I) or frequencies of cytokine-producing cells (J) within the CD4^+^ population. Flow cytometry data is shown as mean + SEM of 12 (Trpm7^+/+^) or 3 (*Trpm7^R/R^*) sick mice per group pooled from 2-3 independent experiments. *p < 0.05, **p < 0.01, unpaired t-test. (**K**) Frequency of CD4^+^ T cells within the CD45^+^ population in the spleen of sick mice. (**L**) Frequency of Vα3.2^+^ transferred T cells within the CD4^+^ population in the spleen. (**M**-**N**) Frequency of FOXP3^+^ (M) or frequencies of cytokine-producing cells (N) within the CD4^+^ population in the spleen. Flow cytometry data is shown as mean + SEM of 12 (Trpm7^+/+^) or 3 (*Trpm7^R/R^*) sick mice per group pooled from 2-3 independent experiments. *p < 0.05, **p < 0.01, ***p < 0.001, ****p > 0.0001 unpaired t-test. (**O**) Representative images of ileal cryosections from Trpm7^+/+^ and *Trpm7^R/R^* mice stained for Vα3.2 (turquoise), CD3e (red) and counterstained with DAPI (blue). Arrows indicate CD3e^+^ Vα3.2^+^ cells. Scale bars = 100 µm for overview images and bar = 20 µm for insets. (**P**) Quantification of the frequency of Vα3.2^+^ within all CD3e^+^ cells. Data is shown as mean ± SEM, dots represent individual mice, *p < 0.05, unpaired t-test. (**Q**) Representative images of ileal cryosections from Trpm7^+/+^ and *Trpm7^R/R^* mice stained for EpCAM (magenta), MHC-II (grey) and counterstained with DAPI (blue). Arrows indicate EpCAM^+^ MHC-II^bright^ cells. Scale bars = 100 µm for overview images and bar = 20 µm for insets. (**R**) Quantification of the median fluorescent intensity (MFI) of MHC-II in EpCAM^+^ cells. Data is shown as mean ± SEM, dots represent individual mice, *p < 0.05, unpaired t-test.

It was shown previously that T_H_17 cells are restimulated in the intestine before migrating to the CNS during transfer EAE (Duc et al., 2019) and we demonstrated previously that gut-homing of T_H_17 cells is impaired in TRPM7 kinase-deficient *Trpm7^R/R^* mice (Romagnani et al., 2017). Thus, we hypothesized that transferred T_H_17 cells may not receive sufficient restimulation in the intestine of *Trpm7^R/R^* mice to cause disease. To test this hypothesis, we performed immunofluorescent staining of gut sections 6 days after T_H_17 cell transfer and prior to EAE onset. Intriguingly, we found significantly reduced numbers of Vα3.2^+^ transferred T_H_17 cells in the ileal lamina propria of *Trpm7^R/R^* recipients as compared to *Trpm7^+/+^* controls (Fig. 4O, P, Fig. S5A, B). Confirming our previous characterization of *Trpm7*^R/R^ mice (Romagnani et al., 2017), this was associated with significantly reduced MHC-II levels on EpCAM^+^ intestinal epithelial cells (IECs) (Fig. 4Q, R, Fig. S5C, D), which may reduce reactivation of Vα3.2^+^ transferred T_H_17 cells in the intestine.

Taken together, loss of TRPM7 kinase activity in host cells leads to reduced disease development in T_H_17-mediated transfer EAE with limited T_H_17 infiltration into the CNS, potentially due to impaired reactivation of the transferred T_H_17 cells in the intestinal lamina propria.

## Discussion

MS is a chronic, demyelinating human autoimmune disease, characterized by the induction of inflammation in the central nervous system (CNS) (Trapp et al., 1999). CNS-invading myelin-specific T_H_17 cells are prime mediators of autoimmune neuroinflammation (Hofstetter et al., 2009). Here, we demonstrate that in CD4^+^ T cells isolated from the cerebrospinal fluid (CSF) of treatment-naïve MS patients, TRPM7 activity was significantly increased compared to CD4^+^ T cells derived from PBMCs of the same patients, which may indicate an enhanced TRPM7 dependency of disease-driving CD4^+^ effector T cells which are enriched in the CSF. Accordingly, our results revealed an upregulation of functional TRPM7 expression not only in acutely activated T cells, but also in nT_H_17 and iT_H_17 cells, with iT_H_17 cells exhibiting the highest surface channel expression per cell, whereas regulatory T cells (T_regs_) showed the lowest number of TRPM7 channels per cell. This further supports an increased dependency of human inflammatory T_H_17 cells on functional TRPM7 expression. These results are in line with recent work, demonstrating the requirement of TRPM7 channel activity for the polarization of naïve human CD4^+^ T cells towards pT_H_17 cells using TGF-ß1, IL-6, IL-21, and IL-23 in the polarizing milieu, with reduced numbers upon pharmacologic inhibition of TRPM7 channel activity, while T_reg_ cell differentiation was unaffected (Madlmayr et al., 2026). Collectively, our CSF patch data indicate that effector T_H_17 cells might be more affected by TRPM7-targeting than T_reg_ cells, making this a potentially suitable therapeutic approach in autoimmune inflammatory contexts such as MS.

T_H_17 cells are characterized by higher plasticity than other T cell subsets and can adopt diverse cell states. On the one hand, pathogenic inflammatory T_H_17 cells are potent drivers of autoimmunity. On the other hand, T_H_17 cells in the intestine are considered “non”-pathogenic and fulfill homeostatic functions in regulation of the gut microbiome. Depending on the differentiating cytokines one can recapitulate the non-pathogenic and pathogenic facets of T_H_17 cells by *in vitro* polarization (Lee et al., 2019; Lee et al., 2012). It should be noted that these cell states are not necessarily final but can still change with changing environments and thus, for example non-pathogenic T_H_17 cells can still develop into pathogenic T_H_17 cells *in vivo* (Hiltensperger et al., 2021; Mangani et al., 2024; Schnell et al., 2021). In the repertoire of MS patients, both pathogenic (pT_H_17) and non-pathogenic T_H_17 (npT_H_17) cells exist (Hiltensperger et al., 2021). Although the initial differentiation of IL-17-producing cells seems to be independent of IL-23, it is expected that IL-23 is required for T_H_17 cells to attain their full pathogenic potential through its maintenance and stabilization of the pT_H_17 phenotype, through suppression of IL-10 production, and through promotion of GM-CSF production and other effector molecules (Lee et al., 2012). Previously, we have demonstrated that the differentiation of *Trpm7^R/R^* naïve T cells into T_H_17 cells without IL-23 is significantly reduced compared to wild-type controls, due to diminished SMAD2 phosphorylation, which is essential for T_H_17 but dispensable for T_reg_ cell polarization (Romagnani et al., 2017). Here we differentiated naïve *Trpm7^R/R^* and WT (Trpm7^+/+^) CD4^+^ T cells in the presence of IL-23 and observed a similar significant reduction in *Trpm7^R/R^* pT_H_17 cells, which was accompanied by a reduced expression of *Rorc*, *Il17a*, *Il17f*, *Il23r*, and *Ccr6*, whereas *Csf2* (GM-CSF) expression was enhanced. This differential behavior of GM-CSF may be due to a different mode of regulation: while most of the pathogenicity-associated factors in T_H_17 cells are mainly induced by elevated RORγt and IL-23R signaling, GM-CSF is additionally regulated by TGF-β signaling, i.e. TGF-β signaling actively represses GM-CSF expression (Codarri et al., 2011; El-Behi et al., 2011). Since we have shown that TGF-β signaling via SMAD2 is diminished in *Trpm7^R/R^* T_H_17 cells (Romagnani et al., 2017), this may explain the increase of GM-CSF in contrast to other pathogenicity-related factors. Similarly, TGF-β is considered to inhibit SOCS3 and thus promote T_H_17 cell differentiation. (Qin et al., 2009) Here our MSigDB hallmark gene set and GSEA enrichment analysis revealed an upregulation of *Socs3* expression in *Trpm7^R/R^* pT_H_17 cells, which could also explain the reduced pT_H_17 cell differentiation. (Qin et al., 2009) Previously, SOCS3 has been linked to MS, as myeloid-specific SOCS3-deficient mice (LysMCre-SOCS3^fl/fl^) were more vulnerable to MOG-induced EAE (Qin et al., 2012). More recently, it was shown that loss of *Socs3* in neutrophils alone (*Socs3*^ΔLy6G^ mice) drives a more severe EAE. (Wang et al., 2026) Another study demonstrated that upregulation of Socs3 expression, due to overexpression of the transcription factor specificity protein 3 (Sp3), which was shown to be downregulated in MS patients (Grekova et al., 1996), reduced CNS inflammation, demyelination and clinical signs in a murine MOG-immunization EAE model. (Li et al., 2025)

While the addition of IL-1β to the T_H_17 polarizing milieu, further increasing the pathogenic phenotype, still resulted in reduced numbers of *Trpm7^R/R^* pT_H_17 cells, transcriptome analysis revealed no significant defects in the expression of T_H_17-associated genes. This might be due to a compensatory effect of IL-1β-induced NFκB signaling, which was shown to be independent of TRPM7 kinase activity (Nadolni et al., 2020). In contrast to other pathogenic T_H_17-signature mRNAs, hallmarks of inflammatory response, including *Ccl3* (MIP1-α), *Ccl4* (MIP1-β), *Ccl5* (RANTES) and *Ccl9* (MIP1-γ) and *Cxcl10* were enhanced in the *Trpm7^R/R^* pT_H_17 cells differentiated without IL-1β. These cytokines/chemokines are considered a uniquely regulated, integral and consistent component of the effector T cell response across different infectious disease models and protective vaccination at the forefront of coordinated host pathogen defenses (Eberlein et al., 2020; Ramesh et al., 2014). In addition, the GSEA enrichment analysis showed a significant upregulation of *Irf1* in several MSigDB hallmark gene sets related to inflammatory responses. The encoded transcription factor IRF1 responds to interferons (Wen et al., 2025) and viral infections, regulating innate and adaptive immunity as well as tumor suppression. (Zoler et al., 2026) Thus, it is tempting to speculate that TRPM7 kinase inhibition would not affect immune responses against diverse pathogens but might be beneficial in the treatment of autoimmune diseases such as MS, particularly as npT_H_17-signature transcripts, including *Foxp3*, *Il6st* (GP130), and *Bach2* are upregulated in *Trpm7^R/R^* pT_H_17 cells. The transcription factor BACH2 (BTB and CNC homology 2) limits immune activation at several stages of CD4^+^ T cell differentiation: it promotes the differentiation of thymus-derived natural regulatory T (nT_reg_) cells, while also restraining effector programs in conventional CD4^+^ T cells, which results in stabilization of peripherally derived iT_reg_ cell differentiation and cell-intrinsic suppression of T_H_1, T_H_2 and T_H_17 cell differentiation (Igarashi et al., 2017). Moreover, large genome-wide association studies have identified single-nucleotide polymorphisms (SNPs) in the *Bach2* locus that are associated with susceptibility to diverse autoimmune and allergic diseases, including MS (Sawcer et al., 2011). Notably, BACH2 is directly phosphorylated by the kinase AKT, which results in impaired nuclear translocation and repressive activity. Thus, the phosphorylation of BACH2 by AKT represents a means by which BACH2 is inactivated (Roychoudhuri et al., 2016; Sidwell and Kallies, 2016). Additionally, it was shown that the repression of BACH2 mediates T_H_17 cell-induced inflammation (Sasikala et al., 2018). We recently demonstrated TRPM7 kinase to directly phosphorylate and thus activate AKT and that naïve T cells require TRPM7 kinase to differentiate into T_H_17 cells (Hoeger et al., 2023; Romagnani et al., 2017). Altogether, these findings suggest TRPM7 kinase as a potential upstream regulator of BACH2. The increased expression of anti-inflammatory genes in *Trpm7^R/R^* compared to *Trpm7^+/+^* pT_H_17 cells prompted us to also investigate potential effects of kinase inactivation on nT_regs_ or the differentiation of naïve T cells into iT_regs_. While the differentiation of iT_regs_ was significantly enhanced upon kinase inactivation, the expression of anti-inflammatory gene signatures was unaltered.

Since inhibition of the TRPM7 kinase reduced pT_H_17 signatures without diminishing the anti-inflammatory profile of T_regs_, we investigated whether kinase inactivation affected the development of EAE utilizing different models. In the conventional MOG/CFA + PTX immunization EAE model, we found no significant effect of TRPM7 kinase inactivation, likely due to the fact that immunization with CFA + PTX induces a strong innate immune response including the release of high levels of IL-1β (Benahmed et al., 2014; Shenderov et al., 2013). Our results demonstrate that the polarization of *Trpm7^R/R^* naïve T cells into pT_H_17 in the presence of IL-1β, although reduced in numbers, restored the diminished expression levels of T_H_17-associated gene signatures in our transcriptome analysis, suggesting that in IL-1β-rich environments it may not be possible to target pT_H_17 cells via TRPM7 kinase inhibition. Mechanistically, we propose a model in which the TRPM7 kinase is required for SMAD2-dependent RORγt and IL-17 signaling but can at least partially be circumvented by the IL-1β-MyD88 pathway (Nadolni et al., 2020). However, in regard to the priming cytokine milieu the MOG/CFA + PTX immunization model is quite artificial: although the exact cytokine milieu present at the time of initial priming of autoreactive T effector cells in MS patients is not known and impossible to study, relapses, which are thought to be associated with reactivation of preexisting and potentially also with priming of novel specificities of autoreactive T cells, have not been described to be associated with high serum levels of IL-1β. Although IL-1β can be detected in the CSF and CNS in a subset of MS patients (Lin and Edelson, 2017), serum levels are lower than in myelin oligodendrocyte glycoprotein antibody associated disease, MOGAD (Kwon et al., 2020), and significantly lower than in autoinflammatory periodic syndromes such as cryopyrin-associated periodic syndrome (CAPS), where serum levels of IL-1β are ∼5fold higher than in healthy controls(Lachmann et al., 2009). Accordingly, while CAPS can be treated well with the IL-1 receptor antagonist Anakinra or the anti-IL-1β antibody Canakinumab (De Benedetti et al., 2018; Rigante et al., 2014), these therapeutic approaches are not currently approved for MS (Hauser and Cree, 2020). Thus, unless CAPS and MS occur in the same patient, which has been described to happen (Schuh et al., 2015), in ‘regular’ MS IL-1β is probably not a disease-defining component of the cytokine milieu. To better model the patient situation, we therefore utilized another EAE model, where disease is induced by transfer of *in vitro* differentiated MOG-specific pT_H_17 cells into wild type recipients in an adjuvant-free setting (Jager et al., 2009). This model mimics the situation of newly diagnosed or relapsing MS patients, where preexisting autoreactive effector T cells are being reactivated during a clinical relapse (Afzali and Korn, 2025; Bielekova et al., 2000; Jelcic et al., 2018). In this pT_H_17 adoptive transfer EAE model, we observed significantly reduced EAE incidence and severity in TRPM7 kinase-deficient recipients, which may reflect a lack of inflammatory signatures in the *Trpm7^R/R^* hosts. While significantly less CD4^+^ T cells infiltrated the CNS in sick *Trpm7^R/R^* mice, especially the frequency of transferred pT_H_17 cells (Vα3.2^+^) was massively reduced, but the frequency of endogenous FOXP3^+^ T_regs_ was increased. This may explain the reduced disease severity of sick *Trpm7^R/R^* mice. We found that the reduced number of transferred pT_H_17 cells (Vα3.2^+^) was associated with a defect of these cells to efficiently migrate into and get reactivated in the intestine due to reduced levels of MHC II expression on intestinal epithelial cells (IECs) in *Trpm7^R/R^* recipients.

Previously studies have suggested that IECs function as noncanonical APCs (Kambayashi and Laufer, 2014), however, the functional role of IEC-derived MHC II initiating autoimmune CD4^+^ T cells and driving CNS inflammation remained unclear. Very recently, it was shown that IECs are critical for the induction of encephalitogenic CD4^+^ T cells during EAE and that IECs up-regulate MHC II in response to neuroinflammatory cues, enabling direct antigen presentation that facilitates T_H_17 differentiation with CNS homing potential (Suzuki et al., 2026). These findings showed clinical relevance, as MS patients exhibited increased expression of epithelial MHC II–associated genes and increased numbers of CD4^+^ T cells in the intestine (Suzuki et al., 2026). Our results presented within this study are in line with these recent reports. Together with our previous findings of altered T cell gut homing and MHC II expression levels in *Trpm7^R/R^* mice (Romagnani et al., 2017), they suggest that even when pathogenic T_H_17 cells are already fully differentiated and present, their reactivation can be successfully inhibited by targeting the TRPM7 kinase.

Altogether, our *in vivo* data directly reflect our observations in *ex vivo* differentiation and transcriptome analyses, with reduced pT_H_17-associated signatures, retained T_reg_ potential and lack of T_H_17 cell reactivation upon TRPM7 kinase inactivation. Thus, the TRPM7 kinase emerges as a prime target candidate in the treatment of MS and potentially other autoimmune diseases.

## Methods

### Animals

For this study we used B6.Cg-FoxP3tm2Tch and B6.Cg-FoxP3tm2Tch/J. T7-R KI from JAX Laboratory Strain #006772 B6-Foxp3^EGFP^ (Haribhai et al., 2007), here called *Trpm7^R/R^* mice. In these mice FOXP3 is co-expressed with an eGFP with restriction to the T cell lineage. In addition, the T7-R KI is carrying a single point mutation at exon 33 position 1646 which leads to a loss of kinase activity (Romagnani et al., 2017). 2D2 mice (C57BL/6-Tg(Tcra2D2,Tcrb2D2)1Kuch/J) were purchased from The Jackson Laboratory. Mice were housed and bred under specific pathogen-free conditions in the Core Facility Animal Models at the Biomedical Center of the LMU Munich or at the Walther Straub Institute of Pharmacology and Toxicology, LMU Munich. The mice were bred in single ventilated cages in groups of 2-5 each cage, with 12 h dark-light circle and food and water ad libitum. Mice were used between 9 to 20 weeks but were age and sex-matched for individual experiments. All animal experiments were designed and performed according to regulations of the animal welfare acts and approved by the animal ethics committee of the state of Bavaria (Regierung von Oberbayern) in accordance with European guidelines and ARRIVE guidelines.

### Sex as a biological variable

Our study examined male and female animals, and similar findings are reported for both sexes.

### Isolation of cells from spleens

Spleens were homogenized and centrifuged. Red blood cell lysis was performed by resuspending the cells in ACK-Buffer (150 mM ammonium chloride; 10 mM potassium hydrogen carbonate; 1 mM Triplex III) for 1-2 min at RT. Lysis was stopped by addition of two volumes RPMI 1640 containing 10 % FBS and the cell suspension was filtered through a 100 µm cell strainer. After centrifugation, the cells were resuspended in RPMI 1640 containing 10 % FBS.

### Murine T_reg_ and T_H_17 differentiation

For T cell differentiation, naïve CD4^+^ T cells were isolated from spleen and LN using the Naïve CD4^+^ T Cell Isolation Kit, mouse (Miltenyi Biotec) according to the manufacturer’s instructions. The remaining CD4-negative cell fraction was treated with 25 µg/ml Mitomycin C (Carl Roth) for 30 min at 37 °C or irradiated at 35 Gy and used as APCs. For T_H_17 differentiation, naïve CD4^+^ T cells were seeded with a density of 1.5 – 2 x 10^6^ cells/ml in RPMI medium containing 10 % FBS, 1 % penicillin-streptomycin, 10 mM HEPES (Sigma), 2 mM L-glutamine, 1 % non-essential amino acids, 1 mM sodium pyruvate and 50 µM β-mercaptoethanol) in the presence of 7.5 – 10 x 10^6^ APCs/ml and stimulated with 2,5 µg/ml anti-CD3 (1455-2C11)), 10 µg/ml anti-IL-4 (11B11), 10 µg/ml anti-IFNγ (XMG.1.2, all BioXCell), 30 ng/ml IL-6) and 3 ng/ml TGF-β1 (both BioLegend) for 48 h at 37 °C. Then, the cells were split 1:3 with fresh medium containing 10 ng/ml IL-23 (BioLegend) and 10 µg/ml anti-IFNγ. For T_reg_ differentiation, naïve CD4^+^ T cells were cultured on a plate coated with anti-CD3 at 0.5 µg/ml in the presence of 10 µg/ml anti-IL-4,10 µg/ml anti-IFNγ and 3 ng/ml TGF-β1. After 48 h, the cells were removed from the coated plate and split 1:3 with fresh medium containing 10 ng/ml IL-2 (BioLegend). On day 4 of culture, successful differentiation was tested by intracellular cytokine staining and subsequent flow cytometry. Gating strategies are shown in Supplementary Figure S7 and S8.

### Isolation of mononuclear cells from CNS

Mice were sacrificed at indicated time points and perfused with PBS through the left cardiac ventricle. Brains were removed from the skull using scissors and forceps and spinal cords were flushed out of the vertebral column with PBS using a 21 gauge needle. To further collect the meninges, the vertebrae were opened with scissors and the meninges were collected with fine forceps. Brain and spinal cord were cut into 1 mm^2^ pieces and digested with collagenase D at a final concentration of 3.75 mg/ml and DNase I with a final concentration of 1.25 mg/ml (both Roche Diagnostic) at 37 °C for 30 min. Afterwards the digested CNS was homogenized by mechanical passage through a 100 µm cell strainer and washed once with cold RPMI Medium for 10 min at 300 x g and 4 °C. Lymphocytes were isolated by density centrifugation using Percoll (Sigma Aldrich): the cell pellet was resuspended in 5 ml of a 70 % Percoll solution and carefully overlayed with 5 ml of a 37 % Percoll solution. Samples were centrifuged for 30 min at room temperature at 650 g, with acceleration and break set to 1. The mononuclear cells were removed from the interface between the two Percoll layers. Then, the cells were washed by RPMI Medium with 10 % FBS and centrifuged at 300 x g for 10 min at 4°C. The cell pellet was then resuspended in 500 µl RPMI containing 10 % FBS and stored on ice.

### Experimental autoimmune encephalomyelitis

For active EAE, mice were immunized subcutaneously (s.c) with 100 µg MOG_35-55_ peptide (BioTrend) emulsified in complete Freund’s adjuvant (CFA containing 5 mg/ml *Mycobacterium tuberculosis* H37 RA (Difco)). Additionally, mice received 150 ng pertussis toxin (PTX, List labs) intraperitoneally (i.p.) on days 0 and 2 after immunization.

For T_H_17 adoptive transfer EAE, naïve CD4^+^ T cells were differentiated into T_H_17 cells as described above. After confirming successful differentiation on day 4 of the culture, the cells were further cultured in the presence of IL-23 and anti-IFNγ until a resting stage was observed on day 8. Then, the cells were collected and restimulated by culturing at a concentration of 2 x 10^6^ cells/ml on a plate coated with anti-CD3 and anti-CD28 antibodies at a concentration of 2 µg/ml. After 48 h, the cells were collected and washed 3 times with PBS. Then, 4 x 10^6^ IL-17-producing cells were injected i.p. into *Trpm7^+/+^* or *Trpm7^R/R^* recipients.

Animals were monitored daily for clinical signs of EAE according to the following criteria: 0, no disease; 0.5, decreased tail tonus; 1, limp tail; 1.5, limp tail and ataxia; 2, hind limb weakness; 2.5 partial hind limb paralysis; 3, complete hind limb paralysis; 3.5, complete hind limb and partial front limb paralysis; 4, complete front and hind limb paralysis; 5, moribund. Mice reaching a score of 2 were provided with gel pads and food pellets on the bottom of the cage.

### Leucocytes from cerebrospinal fluid and blood of MS patients

CSF samples (n=3 patients) were obtained for routine diagnostic procedures in the outpatient department of the Institute of Clinical Neuroimmunology at the LMU Hospital Munich and freshly processed. Blood was taken as reference material from the same time point, and peripheral blood mononuclear cells were separated by density centrifugation and frozen in FBS and 10 % DMSO. Patient characteristics are summarized in Supplementary Table 1. The human material was obtained in compliance with data protection regulations and with the approval of the local ethics committee of the Ludwig-Maximilians-University Munich (Project number: 21-1098). All participants gave their written informed consent in accordance with the principles of the Declaration of Helsinki.

### Mononuclear cells from whole blood

In this study, healthy volunteers of both sexes were enrolled. Informed consent was obtained from all participants. Peripheral blood of healthy volunteers was obtained by venipuncture. The study was conducted according to the guidelines of the Declaration of Helsinki and, approved by the local ethics boards of the Johannes Kepler University Linz (EK 1064/2022).

### Isolation of primary human T cell subpopulations

Cells were isolated from peripheral blood of healthy donors. PBMCs were isolated by density gradient centrifugation using Lymphoprep (Stemcell Technologies, Vancouver, BC, Canada). Isolation of respective lymphocyte subsets was achieved using magnetic cell specific separation kits according to the manufacturer’s protocols. For naïve CD4^+^ T cells, either the Human Naïve CD4^+^ T Cell Isolation Kit II from Stemcell EasySep™ or from Miltenyi Biotec were used, for CD4^+^ T cells, the EasySep™ Human CD4^+^ T Cell Isolation Kit was used. T_H_17 cells were isolated using the EasySep™ Human T_H_17 Cell Enrichment Kit. For both CD4^+^ CD25^-^ effector cells and CD4^+^ CD25^+^ T_reg_ cells, either Stemcell EasySep™ Human CD4^+^CD127lowCD25^+^ Regulatory T Cell Isolation Kit or Miltenyi Biotec CD4^+^ CD25^+^ Regulatory T Cell Isolation Kit, human were used, according to the manuals. A minimum of two different donors were used in primary human T cell experiments.

### *In vitro* human T cell differentiation and flow cytometry staining

For iT_H_17 cells, 1-2x10^5^ cells/ml naïve CD4^+^ T cells were seeded into 96-round-bottom-well plates precoated with 5 µg/ml anti-CD3 (Biolegend) in complete RPMI medium containing the following cytokines: 30 ng/ml rhIL-6 (Biolegend), 100 ng/ml rhIL-21 (R&D System), 30 ng/ml rhIL-23 (Biolegend), 5 ng/ml rhTGF-β1 (BioLegend or Immunotools), anti-10 µg/ml human IFN-*γ* (Thermo Fisher) and 10 µg/ml anti-human IL-4 (BioLegend). Cells were incubated for 6 days by exchanging media (containing respective cytokines) every 2-3 days. On day 6 re-stimulation was performed for 4h with 20 ng/mL PMA (Sigma Aldrich) and 1 µg/mL ionomycin, including 5 µg/mL Brefeldin A (Thermo Fisher). Surface staining was performed using the following antibodies: anti-human CD4-VioBlue (Miltenyi REA623), anti-human CD25-PE (BioLegend, BC96), anti-human CCR6-PEVio770 (Milteny REA190), followed by fixation with 2% formaldehyde in PBS and permeabilization (0.1% Saponin, 1% FBS, 0.005% NaN_3_ in PBS). Signature transcription factor was stained using the following antibody: anti-human RORƔt-PE (Milteny REA278). Cells were analyzed using a Guava Easycyte 6-2L flow cytometer (Luminex Corporation, Austin, TX, USA).

For iT_reg_ cells, 1-2x10^5^ cells/ml naïve CD4^+^ T cells were seeded into 24-flat-bottom-well or 96-well plates in complete RPMI supplemented with 10% NHS. For stimulation, Dynabeads Human T-Activator CD3/CD28 (Gibco or Thermo Fisher, Ratio 1:4), 10 ng/ml rhIL-2 (Preprotech or Immunotools), 5 ng/ml TGF-β1 (Biolegend or Immunotools), 100nM ATRA (Sigma Aldrich) were added and incubated for 5-6 days with intermediary medium exchange on day 4. Surface staining was performed using the following antibodies: anti-human CD4-VioBlue (Miltenyi REA623), anti-human CD25-PE (BioLegend, BC96), anti-human CD45RA-APC-Vio770 (Miltenyi, REA562), anti-human CTLA4-BV605 (BioLegend, BNI3), followed by fixation and permeabilization using the “FOXP3 Fix/Perm” buffer (ThermoFisher Scientific). Signature transcription factor was stained using the following antibody: anti-human FOXP3-APC (Miltenyi, REA1253). Cells were analyzed using a Guava Easycyte 6-2L flow cytometer (Luminex Corporation, Austin, TX, USA) or Cytek Northern Lights 3000 (Cytek Bioscience Inc., Fremont, CA, USA). Naïve CD4 T cells were used as gating control. Data were analyzed with FlowJo® software (TreeStar) and gating strategies are shown in Supplementary figures S6a and S7a.

### Flow cytometry and sorting of murine cells

For intracellular cytokine staining, isolated or cultured cells were stimulated with PMA (50 ng/ml), Ionomycin (1 µg/ml) and Brefeldin A (5 µg/ml) for 4h at 37°C. Viability staining was performed using the Zombie Fixable Viability Kits (Biolegend, 423101 or 423107) according to the manufacturer’s instructions. Subsequently, the cells were washed with FACS Buffer (PBS containing 2 % FBS and 0.05 % sodium azide) and incubated with TruStain FcX^TM^ (1:50; BioLegend) for 15 min at 4 °C to avoid unspecific binding of FACS antibodies to Fc receptors. Surface staining was performed using the following fluorescently labeled anti-mouse antibodies: anti-CD11b (M1/70), anti-CD45 (30-F11), anti-CD4 (RM 4-5), anti-CD19 (RA3-6B2) (all BioLegend). Then, the cells were fixed for 20 min at RT with 0.4 % PFA solution. Cells were washed twice with FACS buffer and resuspended for analysis. For intracellular staining, the cells were stained in permeabilization buffer (PBS containing 2 % FBS, 0.05 % sodium azide, 0.1 % saponin (Sigma)) for 30 min at 4 °C using anti-IFNγ (XMG1.2, BioLegend) and anti-IL-17A (eBio17B7, eBioscience) antibodies. Stained cells were washed with permeabilization buffer and FACS buffer and resuspended in FACS buffer. Samples were measured on a BD FACS Calibur, BD LSRFortessa^TM^ or a Beckman Coulter CytoFLEX S flow cytometer and analyzed with FlowJo^®^ software (TreeStar). For cell sorting, the cells were filtered through a 70 µm cell strainer and sorted using a BD FACSAria^TM^ in the Core Facility Flow Cytometry at the Biomedical Center or at the Institute for Cardiovascular Prevention. Gating strategies for EAE experiments are shown in Supplementary figures S8 and S9.

### Affimetrix microarray analysis for transcriptome data

Cells were isolated and differentiated as previously described. For each condition and sample, 3 - 5 x10^6^ cells were homogenized in TRIzol™ Reagent (Invitrogen™, 15596026) and shipped to ATLAS Biolabs GmbH, which performed the microarray expression profiling via Clariom D Mouse arrays. The total mRNA was extracted by the company using standard protocols. Quality and concentration of the RNA was checked to ensure suitability for microarray analysis.

For analysis we used the probe cell intensity data (CEL) by using the Transcriptome Analysis Console (TAC) Software (version 4.0.3) from ThermoFisher Scientific using the Calriom_D_Mouse Library. The statistical analysis was performed by using the factory default settings. We first generated PCA (Principal Component Analysis) for each Analysis Group and then compared the different conditions with each other getting the Avg(log2), Gene-level Fold change and the p-values.

### Gene-set enrichment analysis (GSEA)

The sample data from TAC Software, were ranked by using the following formula: sign(logFoldChange)x -log10(p-value), were sign(FoldChange) settles the direction of the change (downregulated as negative and upregulated as positive) and -log10(p-value) determines the magnitude of ranking. As reference database the Hallmark gene set (mh.all.v2026.1.Mm.symbols.gmt) provided be the Molecular signatures Database (MSigDB) were selected. We were using the GSEA (v4.4.0) to perform the analysis using ranked scores as described previously(Nassiri et al., 2021).

### Whole-cell patch clamp

The different human T cell populations and the CSF cells were seeded on poly-D-lysin (PDL) coated coverslips and patch clamp experiments were performed in a tight-seal whole-cell configuration at room temperature. Currents were elicited by a ramp protocol from -100 mV to +100 mV over 50 ms acquired at 0,5Hz and a holding potential of 0 mV. By using the EPC-10 (HEKA, Lambrecht, Germany) and Patch Master (HEKA) the high-resolution currents were acquired and recorded. All voltages were corrected for a liquid junction potential of 10 mV between external and internal solutions. Inward current amplitudes were extracted at -80 mV, outward currents at +80 mV. They were normalized to cell size in picofarad (pF) and plotted against the time in seconds (s). To calculate the amount of open TRPM7 channels per cell, we applied divalent-free (DVF) solution, containing (in mM) 140 NaCl, 2.8 KCl, 10 HEPES–NaOH, 10 Cs-EGTA, 5 Na-EDTA, 11 glucose (pH 7.2, 300 mOsm) (at 300s-400s), to already activated TRPM7 channels, thus relieving the divalent cation block of the channel. Utilizing the known single channel conductance (G) of 39 picosiemens (pS) under these conditions at -80 mV (Kerschbaum and Cahalan, 1999), we calculated the number of open channels per cell applying Ohms law and using the equation G [S] = I [A]/U [V]. Patch pipettes were made of borosilicate glass (Science Products, GB 150TF-8P) using the DMZ Zeitz electrode Puller with a resistance of 2–3.5 MΩ. Standard extracellular solution contained (in mM) 140 NaCl, 3 CaCl_2_, 2.8 KCl, 10 HEPES–NaOH, 11 glucose (pH 7.2, 300 mOsm). For DVF experiments, extracellular solution for initial current activation contained (in mM) 140 NaCl, 1 CaCl_2_, 2 mM MgCl_2,_ 2.8 KCl, 10 HEPES–NaOH, 11 glucose (pH 7.2, 300 mOsm) (Romagnani et al., 2017; Zierler et al., 2016). Intracellular solution contained (in mM) 120 Cs-glutamate, 8 NaCl, 10 Cs-EGTA (pH 7.2, 300 mOsm).

### Immunohistochemistry (IHC) of cryosections and image analysis

Mice were sacrificed on day 6 after T_H_17 transfer. The intestine was cut after the stomach and before the anus. Fat was removed and the intestine was cut into 4 parts: duodenum, jejunum, ileum and colon (cecum was discarded). The sections were flushed with cold PBS to remove gut content and cut open longitudinally. Then, the segments were rolled up from proximal to distal with the luminal side inside. Gut rolls were embedded upright in Tissue Tec® O.C.T. Compound (Sakura) and frozen at −80 °C. Cross sections of 10 µm were cut using a Leica CM1950 cryostat. Sections were directly placed on adhesion slides (Superfrost® Plus, Carl Roth) and frozen at −80 °C. For staining, the sections were thawed and fixed in acetone for 10 min and residual O.C.T. compound was removed by washing the sections twice with PBS. Sections were then blocked with PBS containing 1 % BSA and 5 % goat serum for 1 h at RT in a humidified chamber. Subsequently, sections were incubated with the following primary antibodies diluted in PBS containing 1 % BSA for 1 hour in a humidified chamber: hamster anti-mouse CD3e (BD Pharmingen, 1:400), rat anti-mouse B220

(BD Pharmingen, 1:200), rat anti-mouse MHC Class II (I-A/I-E) (BioLegend, 1:200) and biotinylated anti-mouse Vα 3.2 (Miltenyi Biotec, 1:200). Sections were washed three times with PBS containing 0.1 % Tween-20 (PBST) for 7 min and incubated with Alexa Fluor® 568-conjugated anti-hamster IgG (1:400), Alexa Fluor® 633-conjugated anti-rat IgG (1:200) secondary antibodies and Alexa Fluor® 488-conjugated streptavidin (1:400)(all Life technologies) for 1 hour at RT in a humidified chamber in the dark. Then, the sections were washed with PBST three times for 7 min each. If applicable, the sections were stained with Alexa Fluor® 594-conjugated anti-mouse CD326 (EpCAM) (BioLegend, 1:100) in PBS containing 1 % BSA overnight at 4 °C and subsequently washed again three times for 7 min each with PBST. Sections were counterstained with DAPI (Thermo Fisher Scientific) at 1:2000 in PBS for 10 min. The slides were mounted with ProLong Gold Antifade Reagent (Invitrogen) and analyzed in the Core Facility Bioimaging of the Biomedical Center with a Leica SP8X WLL microscope.

Image analysis and quantification was performed using QuPath software(Bankhead et al., 2017): Individual cells were identified using StarDist, a trained model for nuclear segmentation(Schmidt et al., 2018). Cells were classified as CD3e^+^, Vα3.2^+^ CD3e^+^ or EpCAM^+^ based on marker expression and mean fluorescence intensity (MFI) of MHC-II in EpCAM^+^ cells was quantified via QuPath.

### Statistical analysis

Statistical analyses were performed using Graphpad Prism version 10.04 or higher. Appropriate statistical tests were selected as indicated in the figure legends. Differences between two groups were determined using two-tailed student’s t-test. The statistical differences between more than two groups were calculated by one-way ANOVA and multiple comparison t-test. Differences between EAE incidence curves were evaluated using Mantel-Cox Log-rank test. Data summary in bar graphs is expressed as mean ± SEM. The asterisks indicate the statistical differences calculated as: * p<0.05; ** p<0.01; *** p< 0.001 and **** p< 0.0001.

## Acknowledgments

We thank the Core Facility Flow Cytometry and the Core Facility Bioimaging at the Biomedical Center, the mouse facility at the Walther Straub Institute of Pharmacology and Toxicology at the LMU Munich as well as the Core Facility Flow Cytometry at JKU Linz, for providing equipment, services, and expertise. We also thank Dr. med. Habil. Michael Hristov for providing equipment, services, and expertise, Dr. René Platzer for providing expertise in immunology and Michelle Duggan for excellent technical assistance.

This work was supported by the Deutsche Forschungsgemeinschaft (DFG) CRC TRR152, ID 239283807, project P27 (A.P.) and P14 (S.Z.) and by the Emmy Noether Programme PE-2681/1–1, ID 350913910 (A.P.) as well as the JKU Linz, Impetus program ID I-18-23 (BKH).

## Author contribution

KJ and AST conducted animal experiments. KJ, AST, AM designed and conducted experiments, analyzed and interpreted data and edited the manuscript. DL, NRM, KH conducted experiments and analyzed data. LAG performed experiments. FB analyzed and interpreted data. CW, HSK, and TG contributed resources. KJ and SZ prepared figures and wrote manuscript draft. BKH interpreted data, provided supervision and funding and edited the manuscript. AP and SZ conceptualized the study, supervised the project and edited the manuscript. All authors approved the final version of the manuscript. AP und SZ are responsible for the integrity of the work as a whole.

## Competing interests

The authors declare no competing interest

## Data availability

Data are available by the corresponding authors, upon reasonable request.

## Notes

### Competing Interest Statement

The authors have declared no competing interest.

