## Supplemental Figures for "Pathogenicity of autoreactive T_H_17 cells is modulated by the TRPM7 kinase"

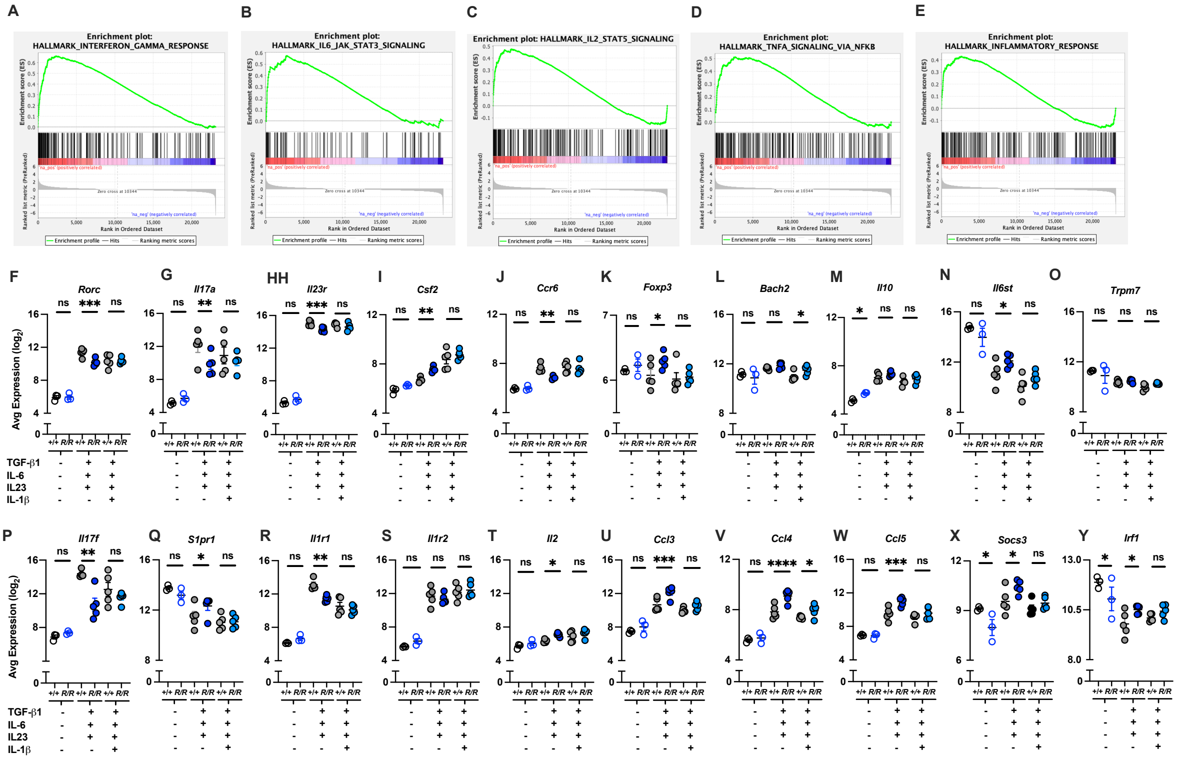


**Supplementary Figure 1. Genetic TRPM7 kinase inactivation inhibits murine iTH17 polarization and inflammatory signatures**

**(A-Y)** Additional data from the transcriptome analysis of TH17 cells differentiated *in vitro* from naïve *Trpm7+/+* (+/+) or *Trpm7R/R* (R/R) CD4$^{+}$ T cells in the presence of IL-6, TGF-β1 and IL-23 with or without addition of IL-1β. (A-E) Enrichments plots of hallmark genes related to inflammatory response. As reference database the Hallmark gene set (mh.all.v2026.1.Mm.symbols.gmt) provided by the Molecular signatures Database (MSigDB) were selected. Pre-ranked data were used to run the GSEA program with 1000 permutations for statistical significance estimation. (F-Y) Average gene expression levels (log2) of indicated genes under indicated experimental conditions. Data is shown as mean ± SEM from 3-5 biological replicates per group. *$p$ < 0.05, **$p$ < 0.01, ***$p$< 0.001, ns, not significant, using one-way ANOVA by the TAC software.

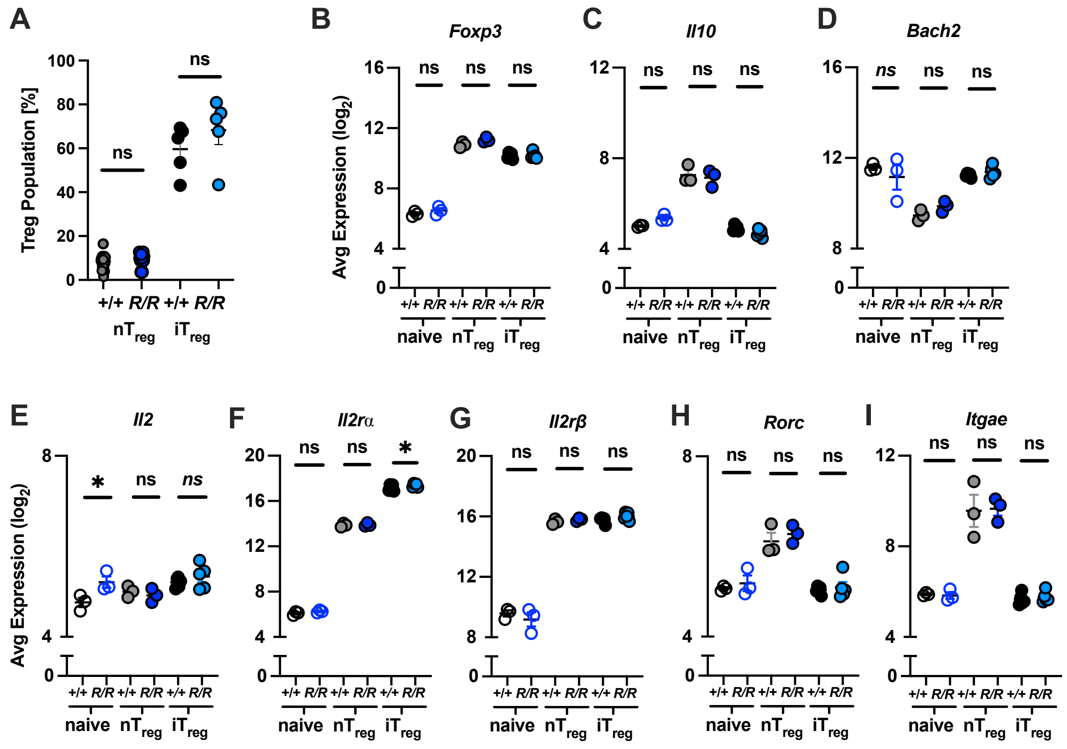


**Supplementary Figure 2. Genetic TRPM7 kinase inactivation does not affect the polarization of murine iT$_{reg}$.** (**A-I**) Transcriptome analysis of nT$_{reg}$ (n= 5), iT$_{reg}$ (n= 5) and naïve T cells (n= 3) from *Trpm7*$^{+/+}$ and *Trpm7*$^{R/R}$ mice. (**A**) Frequency of FOXP3$^{+}$ T$_{regs}$ sorted from splenic CD4$^{+}$ T cells *ex vivo* (nT$_{reg}$) or after *in vitro* differentiation for 4 days (iT$_{reg}$) measured by flow cytometry. Data is shown as mean ± SEM. (**B-I**) Average gene expression levels (log$_2$) of indicated transcripts under indicated experimental conditions. Data is shown as mean ± SEM from 3-5 biological replicates per group *$p$ < 0.05, **$p$ < 0.01, ***$p$< 0.001, ns, not significant, using one-way ANOVA by the TAC software.

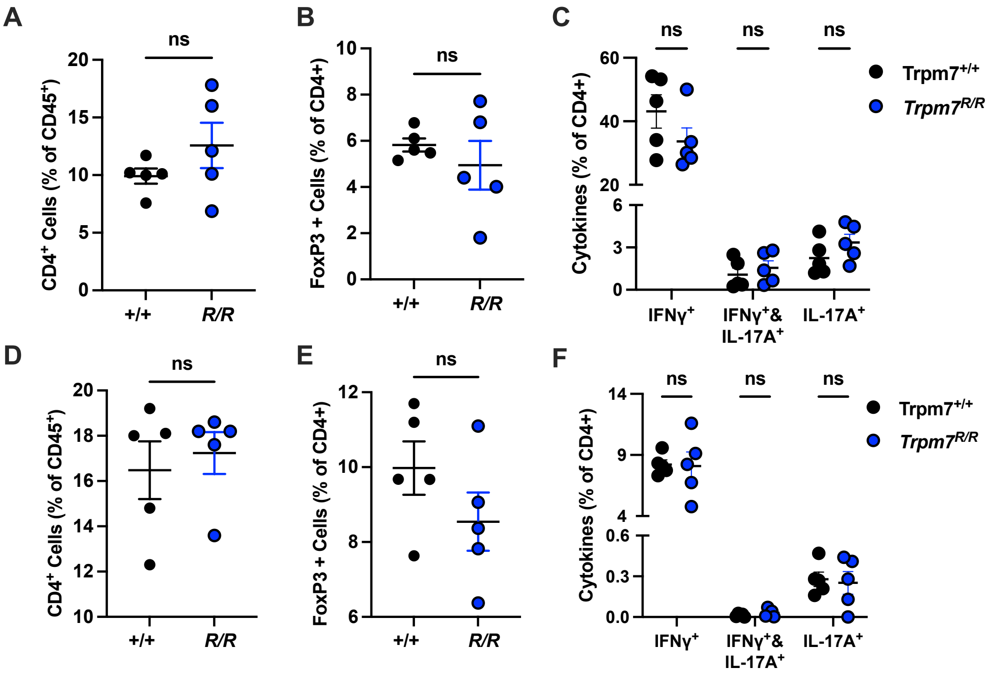


**Supplementary Figure 3. Immune cell status under homeostatic conditions in *Trpm7*$^{+/+}$ and *Trpm7*$^{R/R}$ mice.** (**A**-**F**) Analysis of the distribution of different immune cells (A-C) in the CNS and (D-F) in the SP under homeostatic conditions in Trpm7$^{+/+}$ and Trpm7$^{R/R}$ mice. Flow cytometry data in a-f is shown as mean ± SEM, n= 5 for (*Trpm7*$^{+/+}$) and (*Trpm7*$^{R/R}$) each. ns, not significant.

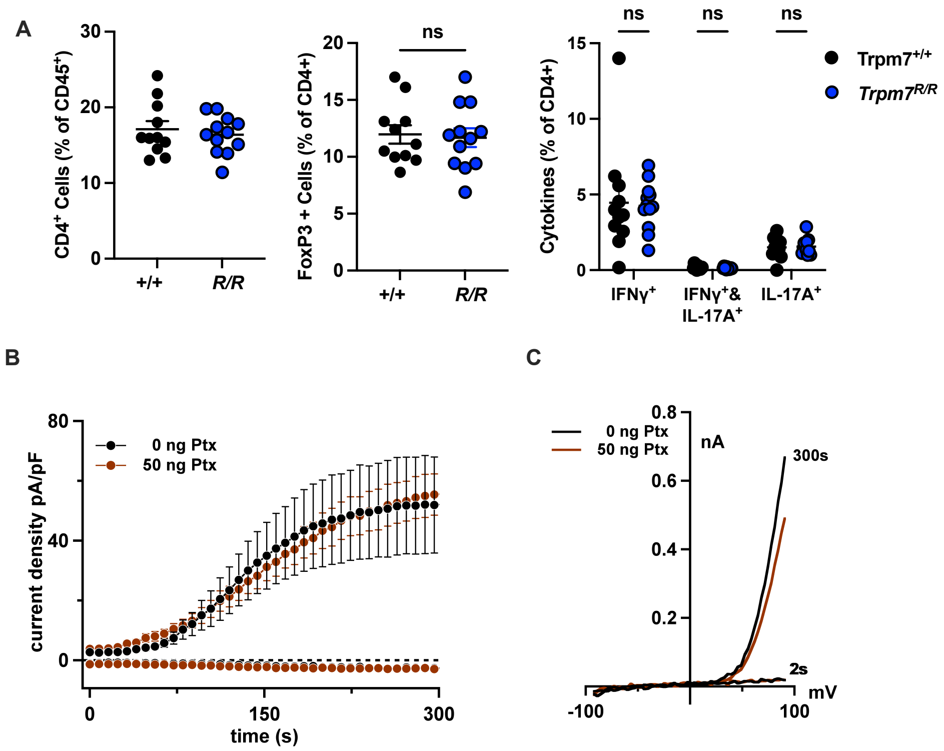


**Supplementary Figure 4. Additional data from MOG/CFA immunization EAE.** (**A**) Frequency of CD4+ T cells within the CD45+ population in the SP of sick mice at the peak of disease. Frequency of FOXP3+ cells and frequencies of different cytokine-producing cells within the CD4+ population in the SP of sick mice at the peak of disease. Flow cytometry data in (A) is shown as mean ± SEM for 11 (*Trpm7*$^{+/+}$) or 12 sick mice (*Trpm7*$^{R/R}$) per group. ns, not significant. (**B**-**C**) Electrophysiological measurements of Jurkat cells, incubated for 4 - 6 h in culture medium (0 ng, black, n= 3) or 50 ng/ml PTX in medium (brown, n=7). (B) Average TRPM7 current density, extracted at +80 mV was blotted *versus* time of the experiment in seconds (s) and (C) average current/voltage relationships (I/V) of Jurkat cells, incubated for 4 - 6 h in culture medium (0 ng, black, n = 3) or 50 ng/ml PTX in medium (brown, n=7) extracted at 300 s. Data are presented as mean ±SEM.

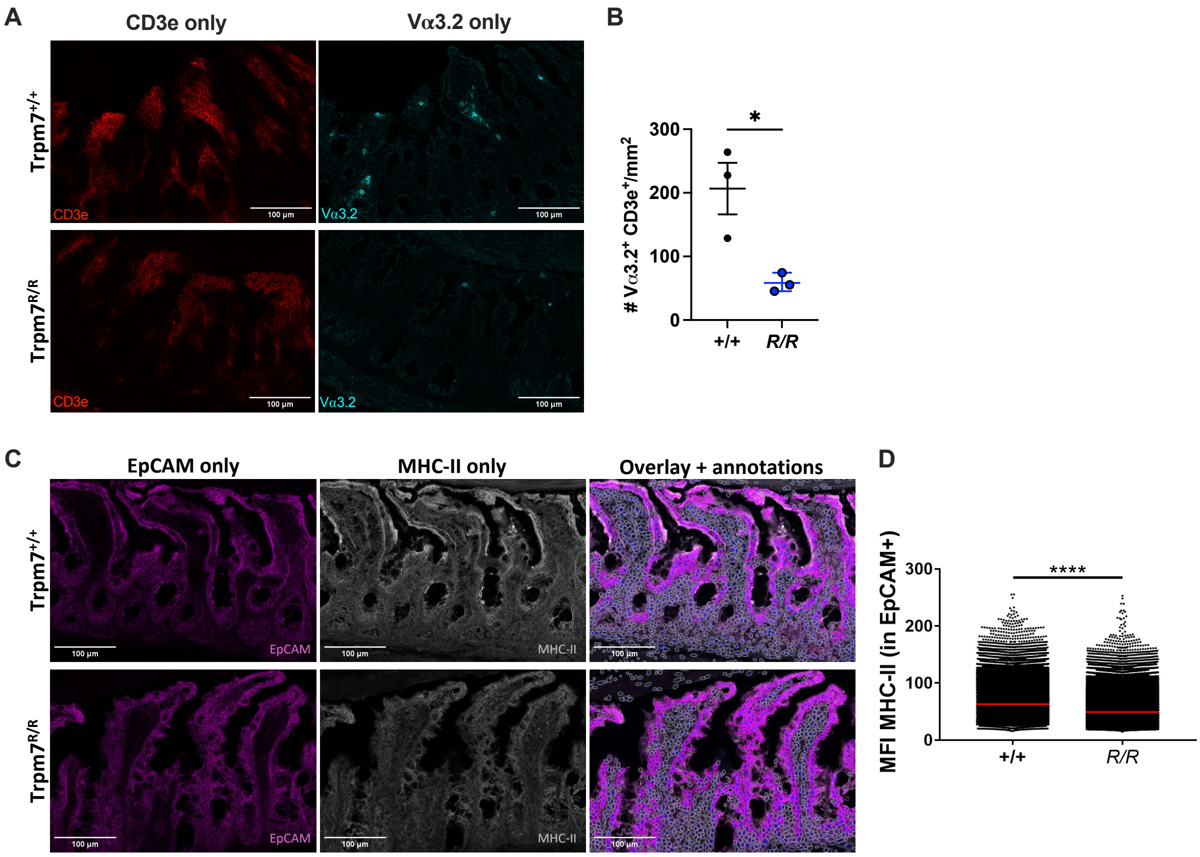


**Supplementary Figure 5. Additional data for IHC staining of intestinal sections from mice**. Representative single stained immunohistochemical visualization of ileal sections of *Trpm7*$^{+/+}$ (top, n=3) and *Trpm7*$^{R/R}$ (bottom, n=3) T$_{H}$17 recipients 6 days post transfer, of (**A**) CD3e (red) and Vα3.2 (turquoise) and (**C**) EpCAM (purple), MHCII (white) and an overlay of both with annotation. (**B**) Quantification of the number of Vα3.2$^{+}$CD3e$^{+}$ cells per mm$^{2}$ in intestinal sections. Dots represent individual mice. (**D**) Quantification of the mean fluorescence intensity of MHC-II in all EpCAM$^{+}$ cells in intestinal sections. Dots represent cells. *p < 0.05, **p < 0.01, ***p < 0.001, ****p > 0.0001 unpaired t-test.

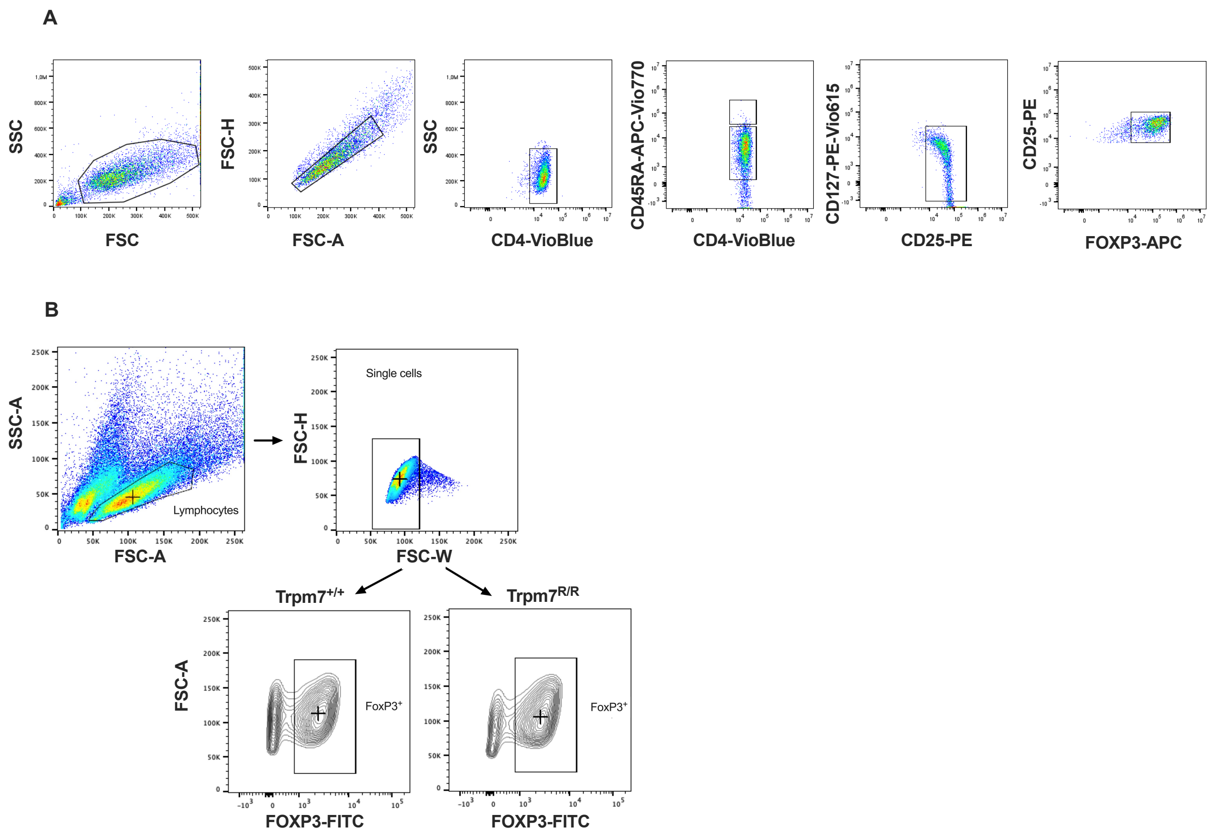


**Supplementary Figure 6. Additional FACS data for $T_{reg}$ populations**

Representative FACS plots and gating strategy of (**A**) human and (**B**) mouse samples.

A

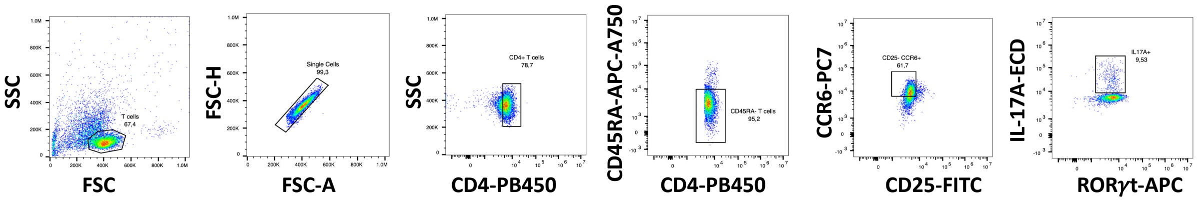


B

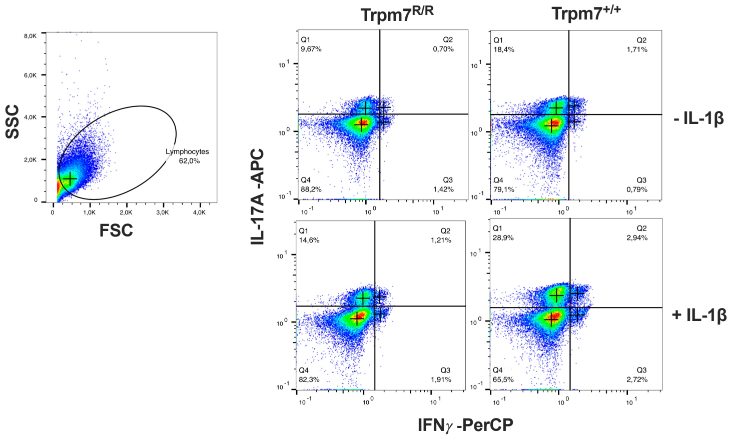


**Supplementary Figure 7. Additional FACS data for $T_H17$ populations** (**A**) Representative FACS plots and gating strategy for human $T_H17$ populations from PBMCs. (**B**) Representative FACS plots and gating strategy for murine $T_H17$ populations shown for *in vitro* differentiated naïve *Trpm7*$^{+/+}$ (+/+) or *Trpm7*$^{R/R}$ (R/R) CD4$^+$ T cells in the presence of IL-6, TGF-β1 and IL-23 with or without addition of IL-1β.

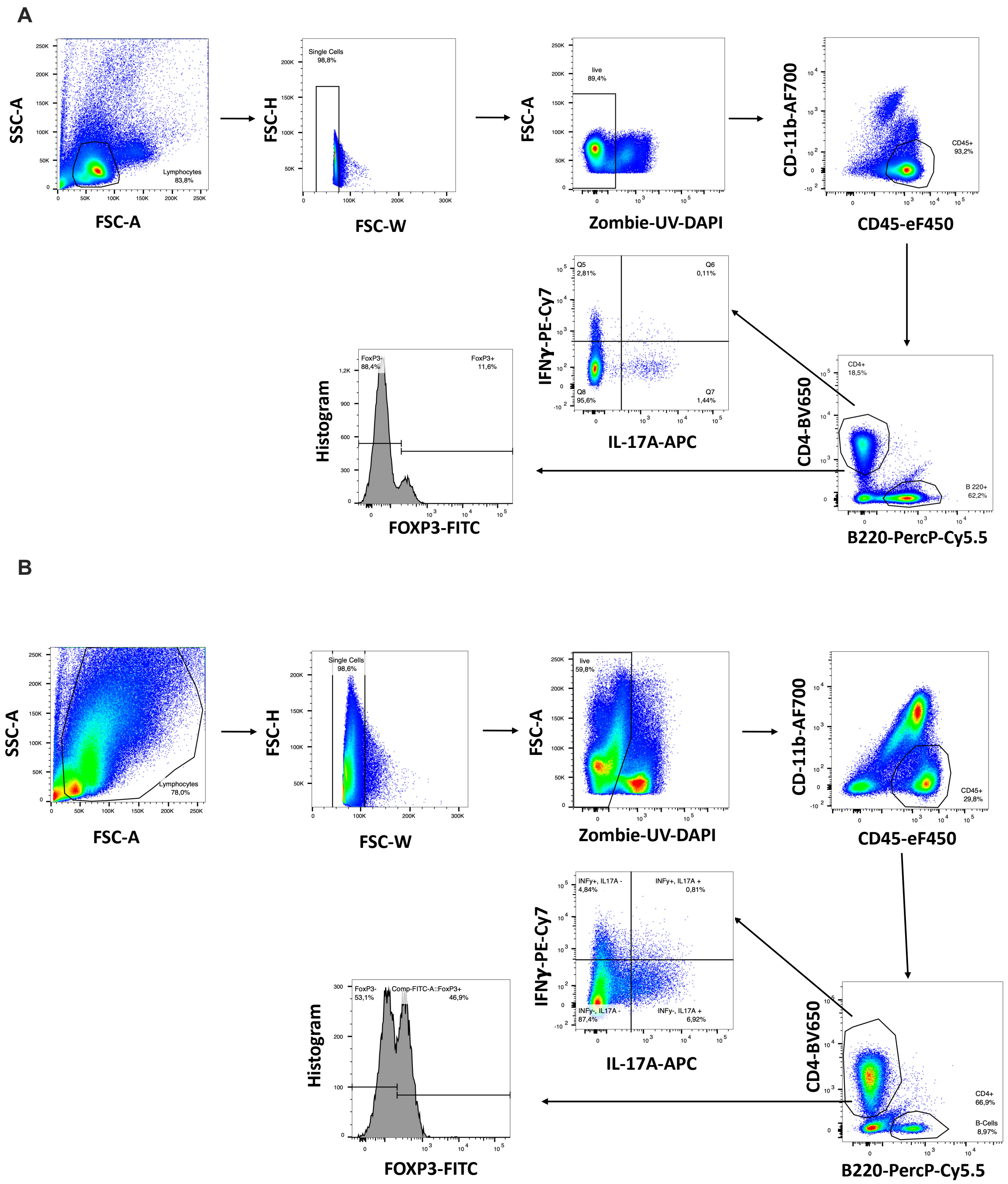


**Supplementary Figure 8. Additional FACS Data for EAE development after immunization with MOG/CFA and PTX.** (**A-B**) Representative FACS plots and gating strategy for (A) spleen and (B) CNS.

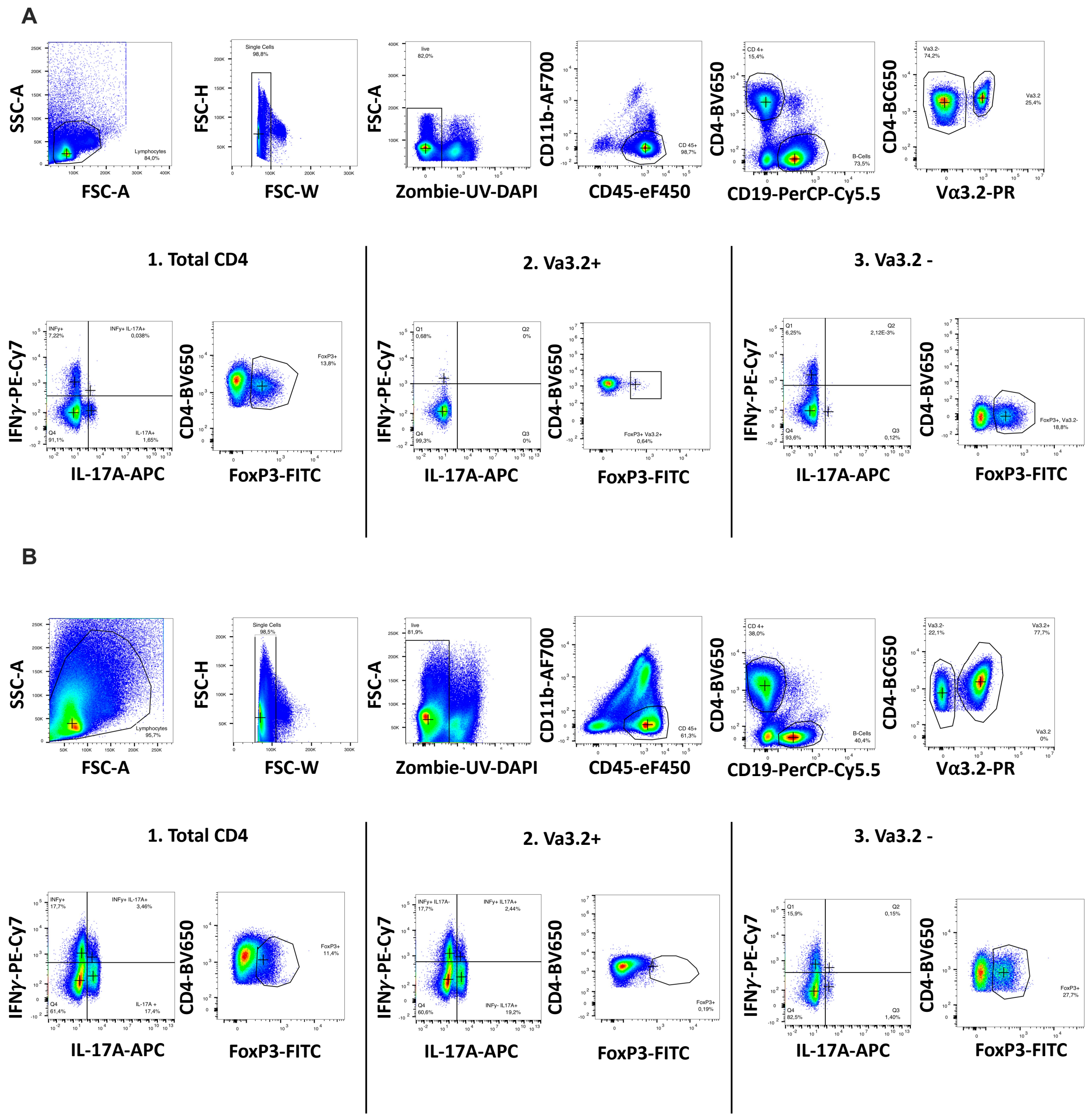


**Supplementary Figure 9. Additional FACS data for $T_H$17-mediated transfer EAE.** Representative FACS plots and gating strategy for (**A**) spleen and (**B**) CNS.

**Supplementary Table 1: Blinded patient data for electrophysiologic analysis.**

| diagnosis | sex | age | disease duration (years) | disease modifying treatment | EDSS | CSF cellcount/µl (Ref. ≤ 5) L | Oliogoclonal bands (TYPE I-IV) |
|---|---|---|---|---|---|---|---|
| RRMS | f | 31 | 0.49 | none | 1.0 | 8 | positive (Type II) |
| RRMS | f | 34 | 3.46 | none | 1.0 | 3 | positive (Type II) |
| RRMS | f | 49 | 14.13 | none | 0 | 3 | suspected (Type II) |

RRMS= relapsing remitting multipls sclerosis
(Type I) OCB negative in Serum and CSF
(Type II) OCB negative in Serum, positive in CSF
(Type III) OCB positive in Serum and CSF, but CSF specific bands present
(Type IV) OCB positive in Serum and CSF, identical bands)
